# Identification of substrates of the protein tyrosine phosphatase PTP1B using *in situ* site-specific photo-crosslinking

**DOI:** 10.64898/2026.07.06.736850

**Authors:** Andrew C. Johns, Yethmie S. Goonatilleke, David C. Cabanero, Yanzhe Ma, Minhee Lee, Anne E. van Vlimmeren, Marko Jovanovic, Neel H. Shah

## Abstract

Protein tyrosine phosphorylation is critical for cellular function, and aberrant phosphorylation is tied to a wide range of human diseases. Identifying the substrates of protein tyrosine phosphatases, the enzymes that erase this modification, is critical to understanding human biology and disease states. The state-of-the-art method for tyrosine phosphatase substrate identification requires the use of mutations that modestly increase the lifetime of enzyme-substrate complexes by kill catalytic activity. While these “substrate-trapping” mutants are useful tools, they work best for high-affinity or abundant substrates that remain phosphatase-bound through cell lysis and enrichment. Here, we use site-specific photo-crosslinking to covalently capture the substrates of tyrosine phosphatases *in situ*. We identify eight different positions around the active site of the phosphatase PTP1B where photo-crosslinker amino acids can be incorporated via amber codon suppression without dramatically disrupting catalytic activity. We then conduct photo-crosslinking experiments in mammalian cells and identify crosslinked proteins by mass spectrometry proteomics, revealing that our approach can capture known PTP1B interactors and substrates. We then show that PTP1B photo-crosslinking *in situ* is sensitive to enzyme localization and identify new PTP1B substrates that regulate contacts between the endoplasmic reticulum and plasma membrane. We also demonstrate that photo-crosslinking can capture signal-dependent interactions. For example, we observe PTP1B crosslinking to the epidermal growth factor (EGF) receptor, a known substrate, in an EGF-dependent manner, and we identify other potential EGF-dependent substrates. Overall, our approach reveals previously unknown roles of PTP1B in signaling systems and could be readily extended to other tyrosine phosphatases in the same family.

## Introduction

Protein tyrosine phosphorylation plays a critical role in signal transduction within cells and is tightly regulated to prevent disease states^1^. Protein tyrosine kinases and protein tyrosine phosphatases (PTPs) work in concert to regulate phosphorylation levels of specific targets in the proteome to maintain homeostasis. There are approximately 90 tyrosine kinases and 100 tyrosine phosphatases in the human proteome^2,3^, which modulate the phosphorylation states of over 10,000 tyrosine phosphosites^4,5^. The vast majority of these phosphosites have not been connected to a specific tyrosine kinase or phosphatase^6^. Importantly, dysregulation in the phosphorylation states of these substrates leads to many disease states such as cancer, immune diseases, and metabolic disorders, among other pathologies^7–9^. Consequently, both tyrosine kinases and tyrosine phosphatases have been the subject of many drug discovery campaigns. While there are around 100 clinically approved tyrosine kinase inhibitors, there are no clinically approved tyrosine phosphatase inhibitors^10–12^. The dearth of clinically approved inhibitors for tyrosine phosphatases likely has two origins: first, tyrosine phosphatases are inherently less “druggable” than tyrosine kinases, due to their lack of an easily-targetable inhibitor binding site^10^, and second, we know less about the signaling functions of most tyrosine phosphatases, including their direct substrates in the phosphoproteome^13,14^. Identifying the specific substrates that are regulated by tyrosine phosphatases is critical to understanding human biology and disease and will likely motivate future therapeutic strategies.

While there have been many tools developed to identify the substrates of tyrosine kinases, we do not have analogous tools to identify the substrates of tyrosine phosphatases due to differences in active site chemistry. For instance, there are highly specific inhibitors and chemical probes for many tyrosine kinases^12^, but relatively few exist for any tyrosine phosphatases^10^, thereby limiting chemical genetics approaches to dissect tyrosine phosphatase signaling. In the same vein, bump-hole approaches have been developed for tyrosine kinases that take advantage of their conserved ATP-binding pocket by making mutations at this site in a target kinase that allow bulky ATP analogues to bind that cannot bind to other unmodified tyrosine kinases^15^. Having a selective ATP co-factor enables labeling of direct substrates for the kinase of interest. By contrast, tyrosine phosphatases lack a cofactor binding pocket to exploit for analogous strategies. Additionally, tyrosine kinase activity leads to a gain of signal (introduction of a modification), enabling selective enrichment and robust detection in phosphoproteomics experiments^16^. Detecting a loss of phospho-signal due to tyrosine phosphatase activity is inherently more difficult^17^. Finally, tyrosine phosphatases have very transient interactions with their substrates as they are quickly dephosphorylated, making traditional methods such as immunopurification infeasible (**Figure 1A**). These challenges have hampered the identification of the direct substrates of tyrosine phosphatases.

**Figure 1.**
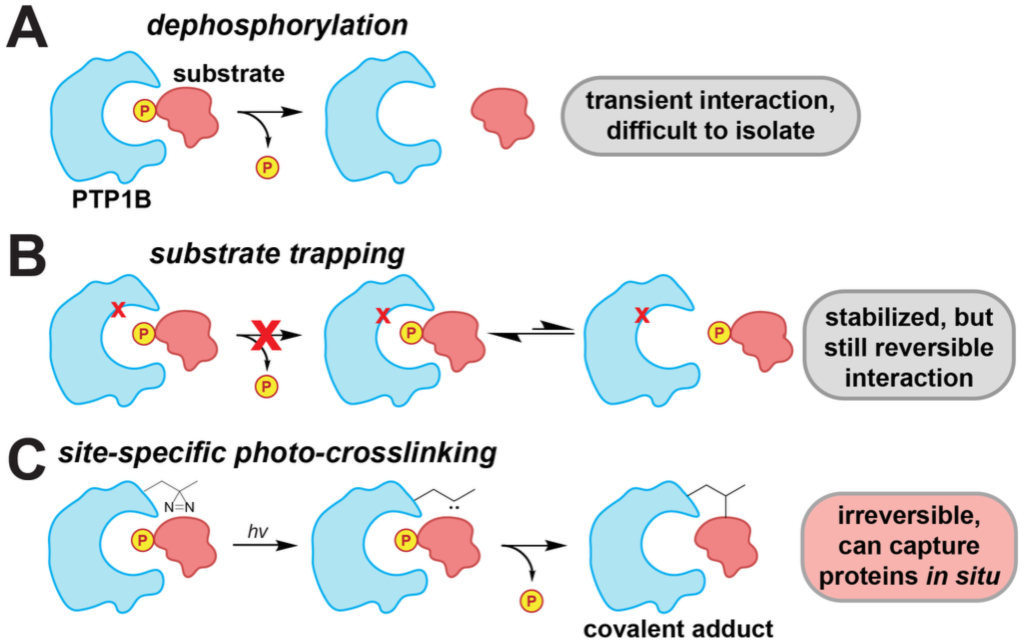
Capturing tyrosine phosphatase substrates by substrate trapping and photo-crosslinking. (**A**) Tyrosine phosphatase substrates are difficult to identify due to their transient interactions with the enzyme. (**B**) Substrate-trapping mutations kill catalytic activity and enhance the lifetime of the enzyme-substrate complex, enabling substrate identification when complexes survive lysis and enrichment. (**C**) Site-specific photo-crosslinking covalently captures transiently interacting substrates *in situ*, allowing for post-lysis enrichment and identification.

The most well-established approach to identify tyrosine phosphatase substrates involves “substrate-trapping” mutations, which kill catalytic activity, thereby increasing the lifetime of enzyme-substrate complexes (**Figure 1B**)^18–20^. While successful in many cases, this approach still requires that the enzyme-substrate complex survives lysis and enrichment to be identified, which precludes the identification of substrates with weak affinity and fast off-rates. Additionally, this approach could potentially capture interactions in the lysate that are not found in the native environment of the cell. To circumvent some of these issues, a recent study combined substrate-trapping mutations with proximity labeling using the promiscuous biotin ligase BioID^21^. In this approach, comparison of proximity-labeling by a PTP-BioID fusion and a substrate trapping PTP-BioID fusion allowed for the identification of proximal proteins that have enhanced biotinylation by substrate-trapping mutants, in a native cellular environment. This method shows significant promise, but it still requires mutations that kill catalytic activity, as well as a fusion to the bulky BioID enzyme. Thus, the field would still benefit from complementary approaches that can facilitate the identification of PTP-substrate interactions and further our understanding of PTP signaling.

In this study, we use site-specific photo-crosslinking to covalently capture and identify substrates of the tyrosine phosphatase PTP1B (encoded by the *PTPN1* gene) (**Figure 1C**). PTP1B is key regulator of a broad range of phosphotyrosine signaling pathways, including insulin and growth factor signaling by receptor tyrosine kinases and cytokine signaling by JAK/STAT proteins, and consequently, it has been the target of several drug discovery campaigns^22^. Furthermore, as the first-discovered member of the PTP family, and given its simple architecture with only one globular domain, PTP1B has served as the central model system for developing PTP-focused tools and understanding structure-function relationships in this enzyme family^23^. Here, using amber codon suppression in mammalian cells, we incorporate photo-crosslinkers at several positions near the substrate-binding cleft of PTP1B. Our crosslinker-equipped variants retain catalytic activity in a cellular context and covalently capture proteins in their native environment, including several known PTP1B substrates and dozens of potential new substrates. Using this strategy, we show that PTP1B dephosphorylates proteins that regulate endoplasmic reticulum-plasma membrane (ER-PM) contacts. Then, using stimulation of the Epidermal Growth Factor Receptor (EGFR) as a case study, we demonstrate that our approach can be used to detect state-dependent changes in the substrate scope of PTP1B. By using site-specific photo-crosslinking of PTP1B, we have expanded the toolkit available to identify substrates of this class of enzymes and have identified new mechanisms of PTP1B signal transduction.

## Results

### Photo-crosslinker-equipped PTP1B constructs retain catalytic activity

Several studies have demonstrated that photo-crosslinker amino acids can be embedded site-specifically into enzymes via amber codon suppression and used to identify direct enzyme-substrate interactions upon irradiation^24–26^. In particular, diazirine-based photo-crosslinkers can be efficiently incorporated into proteins in mammalian cells, and their photo-activation yields a carbene that readily reacts with molecules that are in direct contact^27^. Thus, we reasoned that photo-crosslinking could be a viable strategy to capture direct substrates of PTPs, provided that a crosslinker was strategically placed close to the substrate binding cleft. To identify optimal sites to incorporate a photo-crosslinker into the catalytic domains of PTPs, we first analyzed 29 PTP-phosphopeptide co-crystal structures spanning eight different PTPs and defined a broad substrate binding pocket by identifying residues contacting the substrate peptides (**Figure 2A**). Next, using an alignment of ∼28,000 PTP domain sequences from the Pfam database^28^, we filtered out highly conserved residues that are likely essential for PTP folding or function, and we also excluded glycine residues and positions directly adjacent to catalytic residues (**Figure 2B**). Based on this analysis, we selected eight sites that were proximal to the substrate-binding pocket where substitutions should not perturb PTP function (**Figure 2A,B**). To support this hypothesis, we examined our previously-reported deep mutational scanning data for the isolated PTP domain of a homologous tyrosine phosphatase SHP2^29^, which revealed that the eight chosen sites are highly tolerant of a broad range of amino acid substitutions (**Figure S1**).

**Figure 2.**
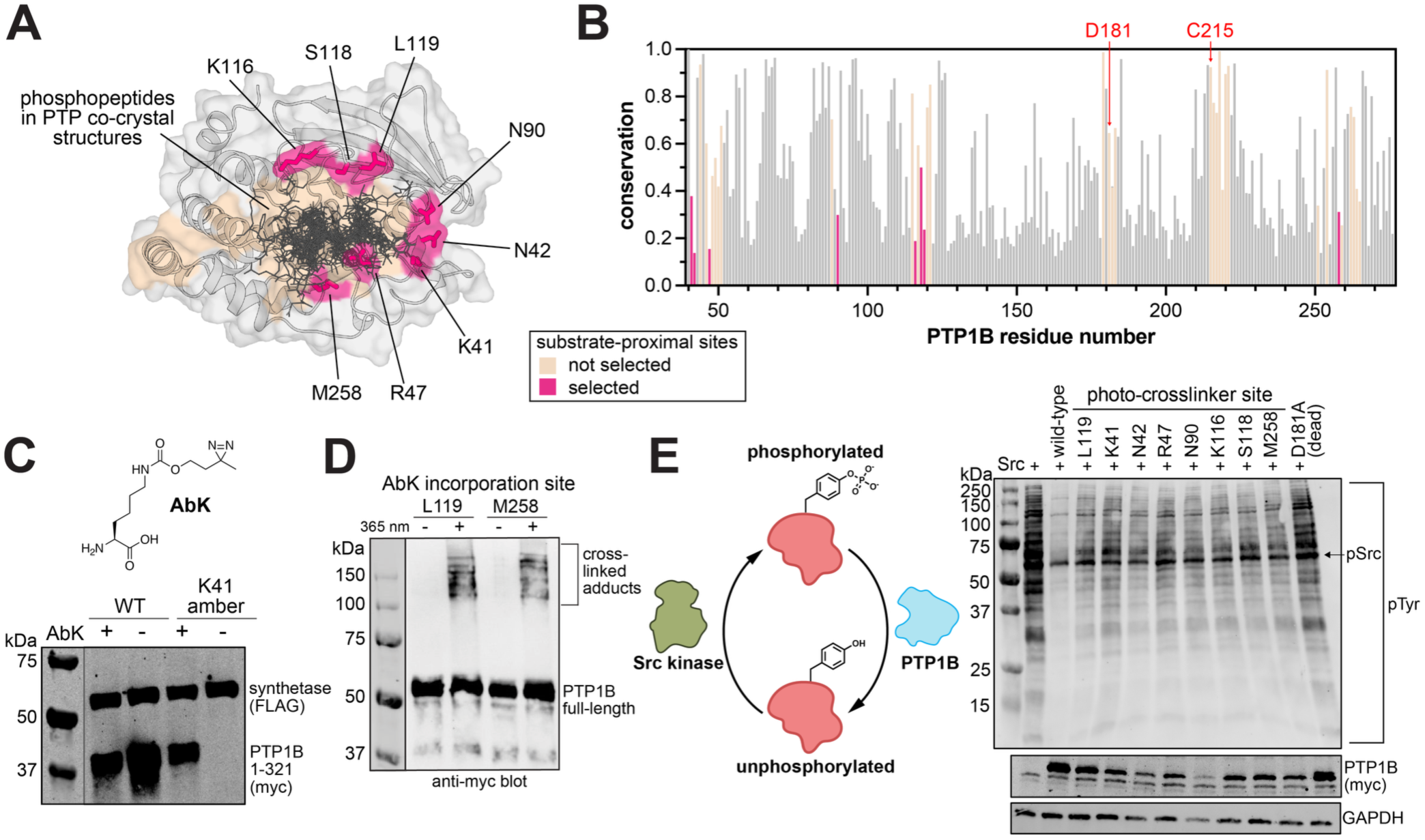
PTP1B-AbK variants retain catalytic activity. (**A**) Crystal structure of PTP1B (PDB code: 1PTU) with phosphopeptides from 29 PTP-substrate co-crystal structures spanning eight different PTPs overlayed. Amino acids in and around substrate binding pocket are colored in tan. Selected photo-crosslinker incorporation sites in pink. (**B**) Conservation scores for each residue in the core of PTP1B, calculated from an alignment of ∼28,000 PTP sequences, obtained from the Pfam database. Conservation is calculated as frequency of the most common amino acid at each position across the alignment. Two important catalytic residues (D181 and C215) are labeled. (**C**) Western blot showing co-expression of the AbK amino-acyl tRNA synthetase (FLAG-tagged), and either wild-type myc-tagged PTP1B (truncated construct, residues 1-321) for a K41-amber variant, in the presence and absence of AbK. (**D**) Western blot showing UV-dependent crosslinking for the full-length PTP1B L119AbK and M258AbK variants. (**E**) Western blot of proteome wide catalytic activity assay of all full-length PTP1B-AbK variants, alongside wild-type PTP1B and the D181A mutant, in the presence of Src^Y529F^.

Next, we evaluated the expression of PTP1B photo-crosslinker variants in mammalian HEK 293 cells via amber codon suppression. We selected the commonly used photo-crosslinker amino acid 3′-azibutyl-N-carbamoyl-lysine (AbK), which has a diazirine moiety that can be converted into a carbene using near-UV (∼365 nm) light^26^. We co-expressed N-terminally myc-tagged PTP1B with an amber codon at position K41, alongside the orthogonal suppressor tRNA and amino acyl tRNA synthetase required for AbK incorporation^25^. Expression of PTP1B from the amber-containing construct was only observed when AbK was added to the media, confirming amber suppression and AbK incorporation (**Figure 2C**). We then examined crosslinking by irradiating cells expressing PTP1B-AbK variants with 365 nm light and blotting for PTP1B. Consistent with prior reports using the AbK crosslinker^25^, we observed robust UV-dependent formation of multiple higher molecular weight species, indicative of protein crosslinking to PTP1B (**Figure 2D**). Critically, different PTP1B-AbK variants had different crosslinking band patterns, suggesting that they may be capturing different protein substrates.

We then expanded our analysis to all eight variants, to examine both amber suppression efficiency and phosphatase activity. All eight PTP1B-AbK variants were compared to the wild-type enzyme and a catalytically dead substrate-trapping mutant (D181A). Although there was less expression of the AbK variants relative to the controls, and some variation in amber codon suppression between variants, all variants could be expressed in HEK 293 cells. The catalytic activity of the PTP1B-AbK variants were then assessed by analyzing global phosphotyrosine levels in the presence of a hyperactive mutant of the tyrosine kinase Src (Src^Y529F^, mouse ortholog). In cells expressing Src alone, we observed substantial tyrosine phosphorylation across the proteome (**Figure 2E**). Co-expression of wild-type PTP1B greatly reduced global phosphotyrosine levels, whereas the catalytically-dead D181A mutant did not impact tyrosine phosphorylation. All eight PTP1B-AbK variants displayed phosphotyrosine levels between the wild-type and catalytically-dead enzymes (**Figure 2E**), demonstrating that these variants retain phosphatase activity.

### PTP1B photo-crosslinker variants have partly overlapping protein capture profiles

Having identified sites for photo-crosslinker incorporation, we profiled all eight full-length PTP1B-AbK variants by photo-crosslinking and proteomics. We expressed our PTP1B-AbK variant of interest, either exposed the cells to 365 nm light or kept them in the dark (on ice), enriched PTP1B using myc-antibody-functionalized beads, then trypsinized the enriched proteins for identification and quantification by mass spectrometry (**Figure 3A**). Triplicate samples with and without UV irradiation were compared to identify photo-crosslinked proteins. In these experiments, we also included wild-type PTP1B as a control, as we found that there was background AbK-independent UV crosslinking (**Figure S2**). A detailed analysis and discussion of background crosslinking can be found in the supplementary text associated with **Figure S2**. Based on our observation of background crosslinking, we focused our analysis on proteins that were enriched by one or more PTP1B-AbK variants but not enriched by the wild-type protein. For each PTP1B-AbK variant, we observed hundreds of proteins enriched >2-fold upon UV irradiation, with dozens of proteins statistically significantly enriched (>2-fold, p<0.05), which we refer to as our low-and high-stringency cutoffs, respectively (**Figure 3B,C and Table S1**). Most of these enriched proteins were not enriched by wild-type PTP1B with the same cutoffs. Notably, all eight variants enriched several known PTP1B interactors and substrates (**Figure 3A, Figure S3, and Table S2**).

**Figure 3.**
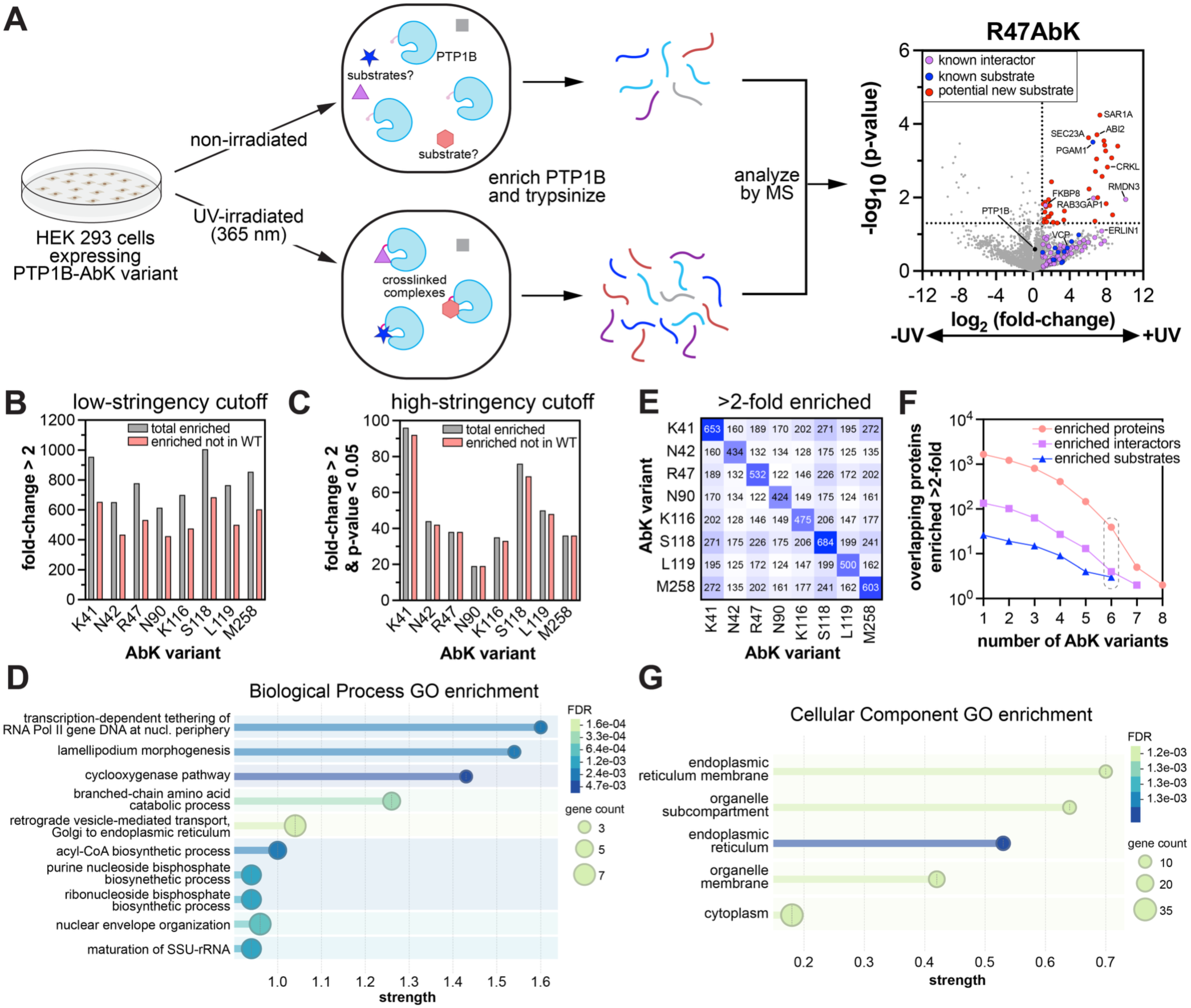
Enrichment of putative PTP1B substrates by site-specific photo-crosslinking. (**A**) Workflow for PTP1B photo-crosslinking and MS proteomics, and volcano plot showing enrichment of proteins by the full-length PTP1B R47AbK variant. Known substrates from DePOD database and literature are blue, known interactors from BioGRID are purple, and putative new substrates are red. (**B**) Comparison of proteins enriched >2-fold upon photo-crosslinking with each PTP1B-AbK variant, excluding those enriched by wild-type PTP1B. (**C**) Comparison of statistically significant (p <0.05) proteins enriched >2-fold upon photo-crosslinking with each PTP1B-AbK variant, excluding those enriched by wild-type PTP1B. (**D**) Biological processes gene ontology analysis of all statistically significant (p <0.05) proteins enriched >2-fold by at least one PTP1B-AbK variant, excluding those enriched >2-fold by wild-type PTP1B. (**E**) Pairwise overlap between PTP1B-AbK proteins, counting proteins enriched >2-fold but excluding proteins enriched by wild-type PTP1B. (**F**) Number of overlapping proteins and interactors enriched >2-fold for a given number of PTP1B-AbK variants, excluding proteins enriched by wild-type PTP1B. (**G**) Cellular component gene ontology analysis of all proteins enriched >2-fold by 6 or more AbK variants, excluding proteins enriched by wild-type PTP1B.

In total, we identified 251 proteins that were statistically significantly enriched (>2-fold, p<0.05) by at least one AbK variant, but not enriched by wild-type PTP1B (>2-fold, with or without a p-value cutoff) (**Table S3A**). Interestingly, a gene ontology (GO) analysis of these proteins identified lamellipodium morphogenesis, which PTP1B is known to modulate^30,31^, as the second most enriched biological process, along with several other biological processes not canonically connected to PTP1B (**Figure 3D**). Several of the 251 significantly enriched proteins were known PTP1B interactors. For example, SRPRα, which was statistically significantly enriched in the K41AbK sample and also enriched by L119AbK, was previously reported to co-localize with PTP1B via proximity-labeling proteomics^32^. SRPRα is a component of the signal recognition particle complex and is responsible for the co-translational insertion of secretory proteins into the ER membrane^33^. SRPRα has one high-frequency tyrosine phosphorylation site (Y261) in the PhosphoSitePlus database^4^, and this residue has previously been shown to be important for mediating protein targeting by the signal recognition particle^34^. Thus, it is plausible that PTP1B may regulate the secretory pathway via direct dephosphorylation of SRPRα. The peptidyl-prolyl isomerase FKBP8, which is found anchored to both the ER and mitochondrial membranes, was statistically-significantly enriched by five PTP1B-AbK variants (**Figure 3A, Figure S3, and Table S3A**). FKBP8 plays a critical role in folding membrane proteins in these two organelles^35^, and it has two prevalent phosphorylation sites (Y322 and Y364) that are not functionally characterized^4^. Using peptide dephosphorylation assays, we validated that FKBP8 pY322 is a high-efficiency substrate for PTP1B, consistent with its strong signal across our photo-crosslinking experiments (**Figure S4**).

While the enriched proteins for individual PTP1B-AbK variants that meet our high-stringency filter (both 2-fold enrichment and p-value <0.05) are likely to be substrates, we also used a complementary strategy to refine a high-confidence list by examining overlap across all eight datasets. Excluding proteins enriched in the wild-type sample, the pairwise overlap between PTP1B-AbK variants ranged from 22% to 45% (**Figure 3E**). Many of the pairwise overlapping enriched proteins were shared across several variants (**Figure 3F**). We identified 38 proteins that were enriched >2-fold by six or more PTP1B-AbK variants but not enriched by wild-type PTP1B (**Table S3B**). 31 of these proteins have at least one tyrosine phosphosite cited on PhosphoSitePlus, and five of these have tandem tyrosine phosphosites (ERLIN1, HCCS, RAB8A, LMAN2, and USP24)^4^. PTP1B has a strong preference for tandem phosphotyrosines, and this motif has been used previously to identify substrates^36–38^. Four of these proteins are also reported interactors in BioGrid^39^ (ERLIN1, SEC20, DNAJC1, and ESYT1), and three are known substrates (PGAM1^40^, VCP^41^, and TTLL12^21^). Notably, several proteins on this list are not documented to be PTP1B interactors or substrates but could plausibly be substrates. For example, Crk like protein (CRKL) was enriched by all 8 PTP1B-AbK variants but not wild-type PTP1B. Its paralog, Crk, is a well-established substrate of PTP1B that is dephoshorylated at the regulatory site Y221^42^. Crk Y221 is known to be a high affinity binding site for its own SH2 domain, and dephosphorylation of this site releases the auto-inhibited state, allowing it to bind phosphotyrosines on other proteins. Given that CRKL is highly homologous to Crk and shares some redundant functions, the analogous site on CRKL (Y207) is likely dephosphorylated by PTP1B^43^. It is also noteworthy that, of the 38 proteins on our high-confidence list, 16 are localized to the endoplasmic reticulum (**Figure 3G**). This suggests that our photo-crosslinking strategy is capturing PTP1B interactions in its native environment, which we explore further in the next section.

In the analyses above, we deliberately excluded proteins that were enriched by wild-type PTP1B upon UV irradiation, however we acknowledge that this approach may filter out some proteins that are *bona fide* substrates but also interact with PTP1B outside of the active site. For example, RMD3 (also known as RMDN3 or PTPIP51) was enriched by seven out of eight AbK variants, but it was also enriched by wild-type PTP1B (**Table S1**). RMD3 is a known interactor of PTP1B^44,45^. Prior studies have shown that RMD3 is phosphorylated by Src at Y176^44^, and phosphorylation of this site is elevated in acute myeloid leukemia cells that have low or no PTP1B expression^46^, further supporting the notion that this site is likely regulated by PTP1B. Given this, the proteins in **Table S3** likely represent a conservative underestimate of PTP1B substrates enriched in our photo-crosslinking experiments.

Finally, although there is overlap between the eight PTP1B-AbK variants (**Figure 3E,F**), several proteins are selectively enriched by one or a small subset of the variants. While this may be due to intrinsic noise in our proteomics experiment, we wondered whether there might be a correlation between crosslinker location and protein capture. However, there was similar level of overlap between pairs of variants, and we could not discern any clear correlation between photo-crosslinker placement and overlap in captured proteins. For example, K41 and N42 are the most proximal photo-crosslinker incorporation sites, with N90 also very close by (**Figure 2A**), but these three PTP1B-AbK variants do not show stronger overlap with one another than they do with other variants (**Figure 3E**). Nonetheless, our global analysis of eight different full-length PTP1B-AbK variants suggests that photo-crosslinkers can be incorporated at a variety of positions around the catalytic site to capture known substrates and discover putative new substrates.

### Photo-crosslinking captures localization-dependent PTP1B substrates

Full-length PTP1B (residues 1-435) is tethered to the endoplasmic reticulum via its C-terminus (**Figure 4A**), but various shorter proteoforms have been identified lacking this tether, including one where the disordered tail is cleaved by the protease calpain^47^, and another bearing just the isolated catalytic module (residues 1-321), which was the first isolated form of PTP1B, from human placenta^48–50^ (**Figure 4B**). We reasoned that photo-crosslinking should allow us to capture differences in PTP1B substrate scope that depend on its localization. Thus, we performed photo-crosslinking proteomics experiments comparing full-length PTP1B to a truncated construct bearing the isolated catalytic module. To focus our analysis, we selected two variants, R47AbK and L119AbK. Both the full-length and truncated PTP1B-AbK variants enriched many known interactors and known substrates, although the truncated constructs had fewer statistically-significant hits (**Figure 4C,D and Table S4**). The slightly higher crosslinking yield for the full-length proteins may be due to increased effective concentration of substrates when the enzyme is confined to the endoplasmic reticulum.

**Figure 4.**
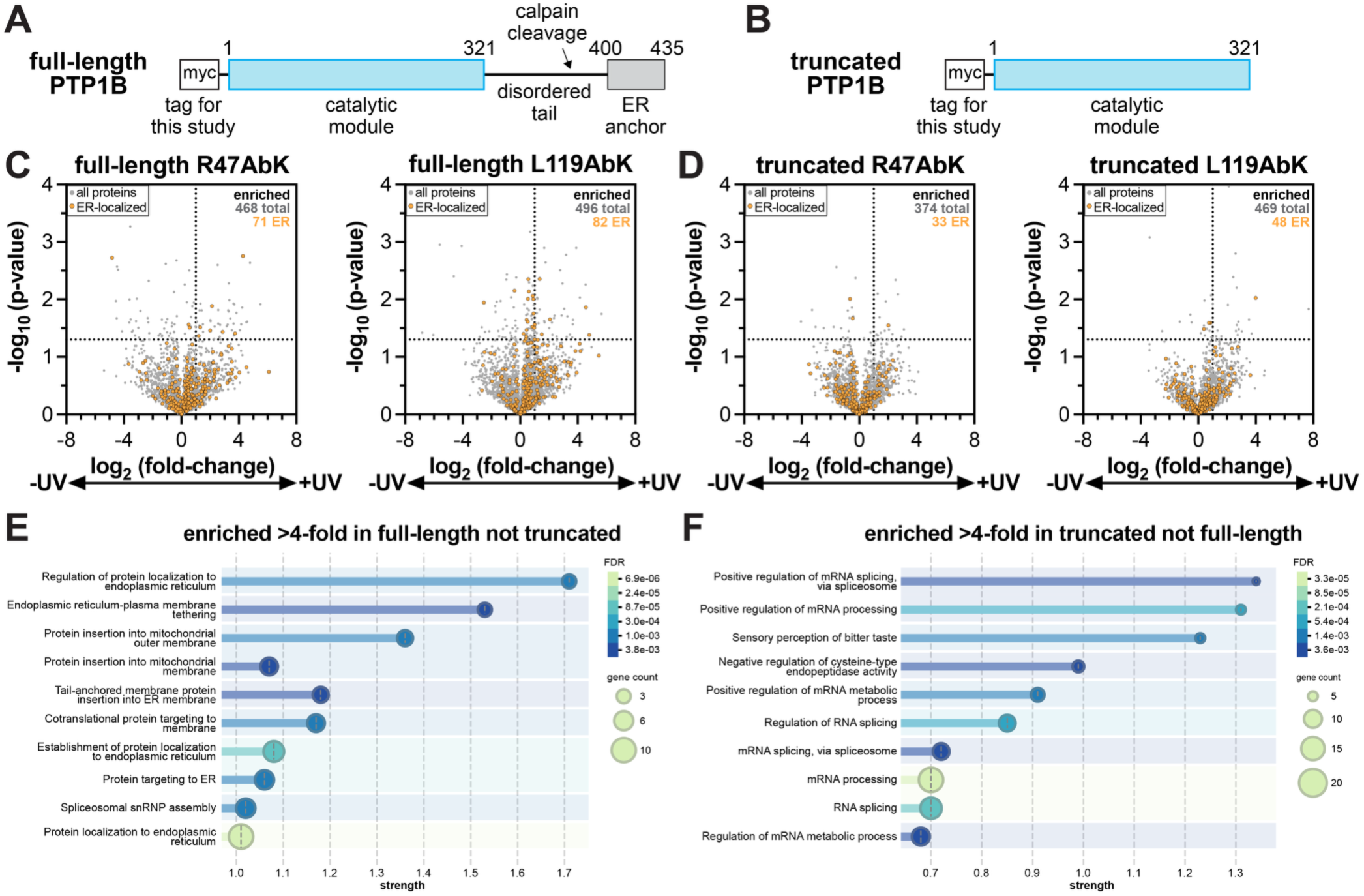
Photo-crosslinking captures localization dependent PTP1B substrates. (**A**) Domain architecture diagram of full-length PTP1B construct. (**B**) Domain architecture diagram of truncated PTP1B construct. (**C**) Volcano plots for photo-crosslinking with PTP1B R47AbK and L119AbK full-length constructs comparing enrichment in +UV and -UV samples. ER-localized proteins are in orange, and other proteins are in gray. The number of total and ER-localized proteins enriched >2-fold is given for each graph. (**D**) Volcano plots for photo-crosslinking with PTP1B R47AbK and L119AbK truncated constructs comparing enrichment in +UV and -UV samples. ER-localized proteins are in orange, and other proteins are in gray. The number of total and ER-localized proteins enriched >2-fold is given for each graph. (**E**) GO analysis of proteins enriched >4-fold for either full-length PTP1B AbK variant but not enriched >4-fold in the truncated constructs. (**F**) GO analysis of proteins enriched >4-fold for either truncated PTP1B AbK variant but not enriched >4-fold in the full-length constructs.

We then asked if there were differences in the localization of the proteins captured in the context of the full-length protein and the truncated form. We first assessed how many of the enriched proteins for each variant are known to be ER-localized and found that the full-length PTP1B constructs enriched disproportionately more ER proteins than the truncated constructs (**Figure 4C,D**). Next, we performed a gene ontology (GO) analysis of the proteins enriched by the full-length constructs but not the truncated constructs (**Figure 4E**), or the truncated constructs but not the full-length constructs (**Figure 4F**). Consistent with the previous section, photo-crosslinking with the full-length R47AbK and L119AbK variants enriched many proteins involved in ER-related processes, including regulation of protein localization to endoplasmic reticulum, tail-anchored membrane protein insertion into endoplasmic reticulum, and endoplasmic reticulum-plasma membrane tethering (**Figure 4E**). By contrast, this ER signature was not observed for the truncated constructs (**Figure 4F**). These results demonstrate that photo-crosslinking captures protein interactions *in situ* and can preserve information about PTP1B subcellular localization.

### PTP1B dephosphorylates endoplasmic reticulum-plasma membrane tethering proteins

As noted in prior sections, our crosslinking experiments identified several putative PTP1B substrates in the ER. One pair of interesting proteins that emerged from our experiments were Extended Synaptotagmin-1 (ESYT1) and Extended Synaptotagmin-2 (ESYT2). These are ER bound proteins that pinch the ER to the plasma membrane, bringing proteins on both interfaces in close proximity^51^. This is relevant to PTP1B signaling because PTP1B dephosphorylates a wide range of receptor tyrosine kinases in response to extracellular signals such as VEGF, EGF, and insulin, among others^52–55^. This pinching effect could be important in mediating PTP1B access to these receptor tyrosine kinases on the plasma membrane. The structure of ESYT proteins is composed of a synaptotagmin-like mitochondrial binding protein (SMP) domain followed by five C2 domains in ESYT1 or three C2 domains in ESYT2 (**Figure 5A-C**). Calcium binding to the third C2 domain of ESYT1 (C2C) releases its autoinhibited fifth C2 domain (C2E), allowing it to bind phosphatidylinositol 4,5-bisphosphate (PIP2) and bring the ER to the plasma membrane^53^. For ESYT2, PIP2 binding to the third C2 domain (C2C) allows it to pinch the ER-plasma membrane gap^57^.

**Figure 5.**
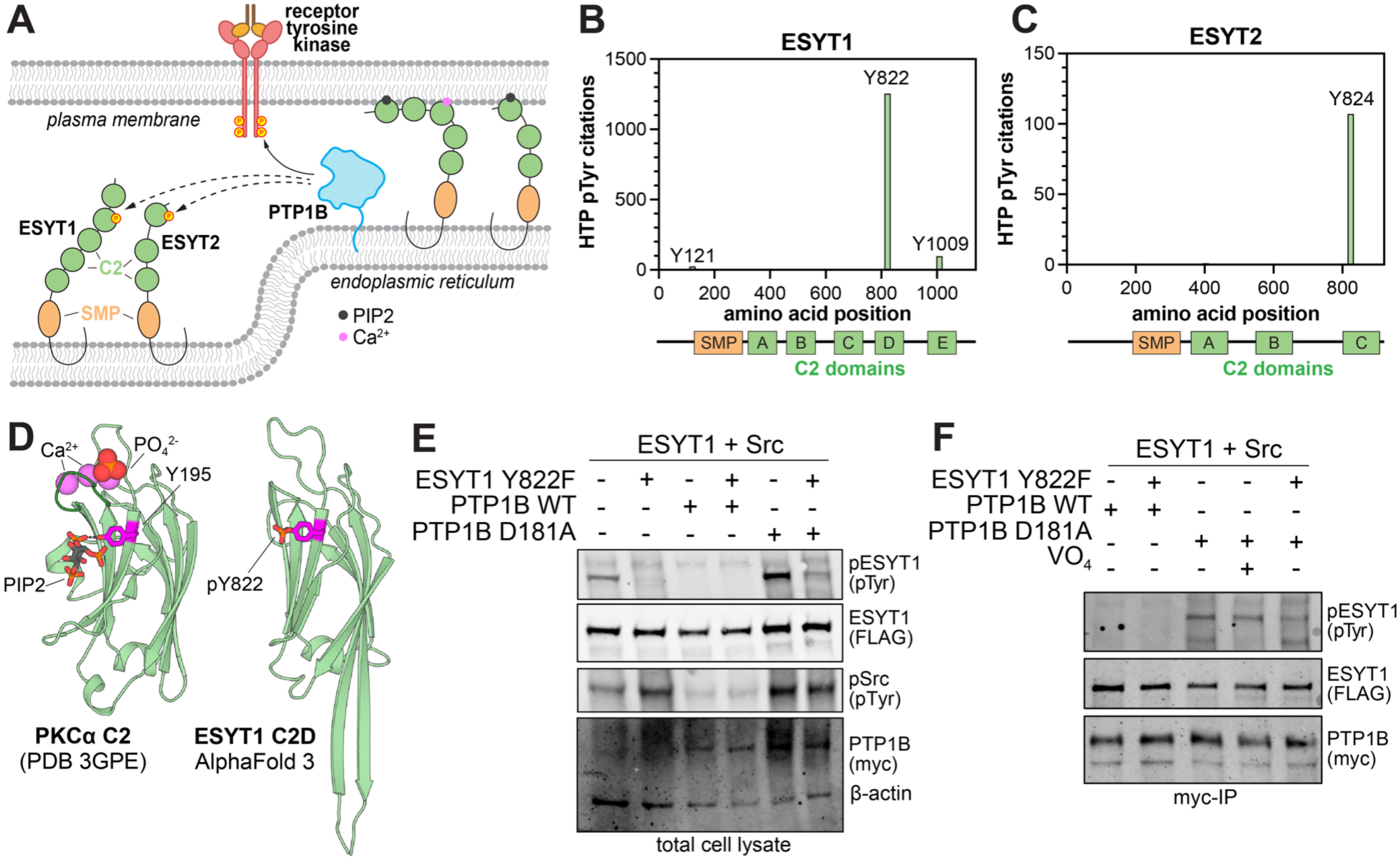
Validation of ESYT1 as a substrate of PTP1B. (**A**) Diagram depicting the pinching effect of ESYT1 and ESYT2, and potential regulation of these proteins by PTP1B. (**B**) High-throughput (HTP) proteomics citations of the tyrosine phosphosites (pTyr) at each position on ESYT1, with the domain diagram of ESYT1 aligned below. (**C**) High-throughput (HTP) proteomics citations of the tyrosine phosphosites (pTyr) at each position on ESYT2, with the domain diagram of ESYT2 aligned below. (**D**) Crystal structure of PKCα C2 domain with calcium, phosphate, and PIP2 bound (PDB code 3GPE), highlighting an interaction between PIP2 and Y195 (analogous to Y822 and Y824 on ESYT1 and ESYT2, respectively). An AlphaFold 3 model of the ESYT1 C2D domain, with phosphorylated Y822, is shown for comparison. (**E**) Total cell lysate western blot of FLAG-ESYT1 wild-type or Y822F co-expressed with Src kinase Y529F and myc-PTP1B WT or D181A variants. (**F**) Myc-PTP1B IP from samples co-expressing ESYT1 wild-type or Y822F mutant and Src. Vanadate (VO_4_) is added as competitive orthosteric inhibitor of PTP1B.

Interestingly, ESYT1 C2D and ESYT2 C2C bear tyrosine phosphorylation sites that are frequently observed in phospho-proteomic datasets (**Figure 5B,C**)^4^. These tyrosines, ESYT1 Y822 and ESYT2 Y824, are at the same position on their respective C2 domains. A published crystal structure of the C2 domain of PKCα shows that this highly-conserved tyrosine residue hydrogen bonds with one of the phosphate groups of PIP2^58^ (**Figure 5D**). Although the phosphorylation sites on ESYT1 and ESYT2 have not been biochemically characterized, presumably their phosphorylation will disrupt PIP2 binding. For ESYT2, this would impede its binding the plasma membrane and disrupt its pinching function. The same may be true of ESYT1, although there are no reports directly describing the functional effects of PIP2 binding to its fourth C2 domain. Thus, if PTP1B dephosphorylates these sites, the phosphatase would be a positive regulator of ESYT1/ESYT2 function.

To confirm that the ESYT1 Y822 phosphosite is a substrate of PTP1B, we first identified a kinase that can sufficiently phosphorylate this site for validation studies. Src was chosen first, because it is activated in response to growth signals and is a substrate of PTP1B, exemplifying how they are often intertwined in cellular pathways^59^. To ensure that regulation of Src by PTP1B does not obfuscate the data, we used the hyperactive Src^Y529F^ variant (mouse numbering), which lacks the key phosphosite on the Src regulatory tail that is typically dephosphorylated by PTP1B^60^. We then co-expressed both wild-type PTP1B and the substrate-trapping variant D181A in the presence of Src^Y529F^. Analysis of the total cell lysate showed that Src^Y529F^ phosphorylated wild-type ESYT1 but not a Y822F mutant, indicating that Y822 is being phosphorylated (**Figure 5E**). Co-expression of wild-type PTP1B dramatically reduced ESYT1 phosphorylation, whereas co-expression of the substrate-trapping variant of PTP1B led to an observed increased wild-type ESYT1 phosphorylation but not that of ESYT1 Y822F (**Figure 5E**). The substrate-trapping variant likely forms a complex with ESYT1 protecting this site from dephosphorylation, thereby increasing phosphorylation levels at this site. In a myc immunopurification from these cell lysates, wild-type and Y822F ESYT1 were pulled down in equal amounts irrespective of phosphorylation state, suggesting that ESYT1 has a secondary binding site on PTP1B beyond binding to the active site as a substrate (**Figure 5F**). Indeed, addition of the orthosteric PTP1B inhibitor orthovanadate during immunopurification did not disrupt the PTP1B-ESYT1 complex, indicating a secondary non-catalytic interaction (**Figure 5F**).

We then set out to validate ESYT2 as a substrate, however, we were unable to sufficiently phosphorylate ESYT2 to enable efficient substrate-trapping experiments. Given our results for ESYT1, we first attempted to phosphorylate ESYT2 by Src co-expression, or by incubation of cell lysates with the Src kinase domain, but we did not observe ESYT2 phosphorylation. We then tried an activated form of the FGFR1 kinase domain because ESYT2 is known to interact with FGFR1^61,62^, but again observed no phosphorylation. As an alternative validation approach, we synthesized phosphopeptides spanning the pY822 and pY824 sites on ESYT1 and ESYT2, respectively, and measured the Michaelis-Menten kinetic parameters for their dephosphorylation by PTP1B. These experiments revealed that, at the peptide level, ESYT2 pY824 is a 6-fold better substrate than ESYT1 pY822, but not as efficient as high-activity PTP1B substrates, such as the EGFR tail phosphosites^63^ (**Figure S4**). The higher activity for the ESYT2 peptide relative to the ESYT1 peptide, coupled with our validation of ESYT1 pY822 as a substrate of PTP1B, suggests that ESYT2 could plausibly also be a substrate of PTP1B. Furthermore, the low activity for the ESYT1 peptide indicates that additional interactions outside of the active site of PTP1B and the local sequence around the phosphosite are likely required for efficient dephosphorylation, consistent with our orthovanadate competition experiments (**Figure 5F**).

We note that there were many other interesting ER proteins that were hits in our photo-crosslinking experiments. Two of these, the cytochrome P450 oxidoreductase (POR) and Endoplasmic Reticulum Raft-associated protein 1 (ERLIN-1), were enriched by seven mutants and are known interactors but not known substrates, although POR was also enriched by wild-type PTP1B. Additionally, these two proteins contain tandem phosphotyrosine sites, which PTP1B prefers, as discussed earlier. Regulation of POR may have broad effects on cellular metabolism, while ERLIN-1 regulates calcium signaling, facilitates the degradation of PIP3, governs cholesterol metabolism, and regulates cell fate by promoting autophagy^64^.

### Photo-crosslinking captures EGF-dependent protein substrates

Lastly, we aimed to examine whether photo-crosslinking could be used to capture signal-dependent changes in the PTP1B substrate repertoire (**Figure 6A**). We used EGFR stimulation by EGF as a case study, because PTP1B is known to modulate EGFR signaling and vesicular trafficking^65,66^, and EGFR itself is a direct substrate of PTP1B once it is activated and auto-phosphorylated^18^. In parallel with EGFR simulation, inspired by a previous study that combined substrate-trapping mutations with proximity-labeling proteomics^21^, we explored whether photo-crosslinking could be combined with substrate trapping (using the D181A mutation) for enhanced substrate capture. To test these ideas, we conducted four photo-crosslinking experiments with cells expressing PTP1B bearing L119AbK: (i) L119AbK in unstimulated cells, (ii) L119AbK in EGF-stimulated cells, (iii) L119AbK+D181A in unstimulated cells, (iv) L119AbK+D181A in EGF-stimulated cells. In each case, cells were either non-irradiated or irradiated with near-UV light, and all conditions were conducted in triplicate. We first analyzed how many proteins were enriched when comparing the +UV to -UV samples and found little variation between conditions (**Figure 6B and Table S5**). We then applied a high-stringency p-value cutoff of 0.05 and found that fewer proteins met this criterion in the samples stimulated with EGF (**Figure 6C and Table S5**). Notably, among the enriched proteins in the +EGF samples was EGFR. Based on this analysis, we concluded that our method likely detects signal-dependent changes in PTP1B substrate scope.

**Figure 6.**
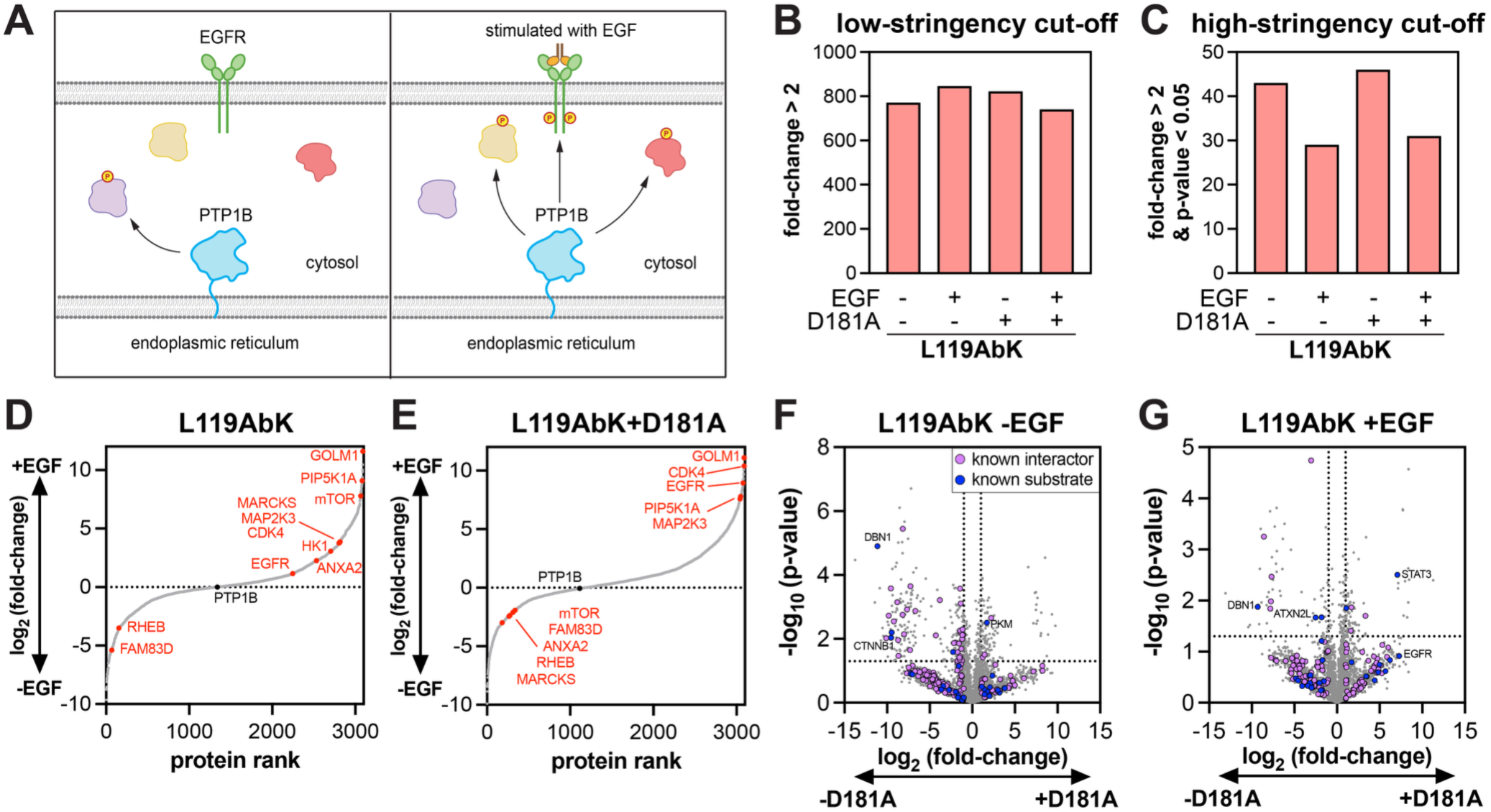
Photo-crosslinking captures EGF dependent substrates. (**A**) EGF-stimulation-dependent changes in PTP1B substrates. (**B**) Proteins enriched >2-fold by photo-crosslinking to the L119AbK variant, +/- addition of a substrate-trapping D181A mutation and +/- stimulation with EGF. (**C**) Proteins statistically significantly (p<0.05) enriched >2-fold by photo-crosslinking to the L119AbK variant, +/- addition of a substrate-trapping D181A mutation and +/- stimulation with EGF. (**D**) Protein rank order for EGF-dependent enrichment after photo-crosslinking with the PTP1B L119AbK variant. Enriched and depleted EGF-signaling-related proteins in red. (**E**) Protein rank order for EGF-dependent enrichment after photo-crosslinking with the PTP1B L119AbK+D181A variant. Enriched and depleted EGFR-signaling-related proteins in red. (**F**) Volcano plot comparing L119AbK and L119AbK+D181A UV-irradiated samples in without simulation by EGF. (**G**) Volcano plot comparing L119AbK and L119AbK+D181A UV-irradiated samples in with stimulation by EGF. In panels F and G, known substrates from DePOD database and literature are blue, and known interactors from BioGRID are purple.

To gain further insights into EGF-dependent changes in PTP1B substrates, we directly compared the UV-irradiated samples +/- EGF to identify the proteins that are enriched or depleted upon stimulation by EGF (**Figure 6D,E and Table S6**). The proteins that are enriched in these comparisons are likely to be phosphorylated in response to EGFR stimulation making them more available for photo-crosslinking to PTP1B. The proteins that are depleted in the +/- EGF comparison could have reduced phosphorylation levels upon EGFR stimulation, making them less available for dephosphorylation and thus less likely to be photo-crosslinked. Reassuringly EGFR is enriched upon stimulation by EGF both for the L119AbK variant and the L119AbK+D181A variant (**Figure 6D,E**). Many other proteins that are not known PTP1B substrates but are directly involved in the EGFR signaling cascade were also enriched. For example, in both the L119AbK and L119AbK+D181A datasets, GOLM1, which has one reported but functionally uncharacterized tyrosine phosphosite (pY351), was the most enriched, statistically significant protein in response to EGFR stimulation. GOLM1 is a Golgi resident protein that mediates EGFR anchoring to the trans-Golgi network and recycling of EGFR back to the plasma membrane for sustained signaling^67^. CDK4 was also found to be highly enriched in our analysis. CDK4 is canonically known to drive cell division by controlling the transition from G1 to S phase^68^. It is also known to be basally phosphorylated at Y17 by Src family kinases during quiescence and dephosphorylated when entering the cell cycle^69,70^.

Another protein that displayed EGF-dependent enhancement in photo-crosslinking to PTP1B was Hexokinase-1 (HK1) (**Figure 6D**). This key metabolic enzyme contains a regulatory site at Y732 that is a high-frequency phosphosite in the PhosphoSitePlus database^4^. Phosphorylation of this site in response to growth signals allows HK1 to dimerize, causing an increase in activity, shifting the cell into glycolysis^71^. Y732 is also basally phosphorylated and its phosphorylation increases in response to growth signals such as EGF^71^. Furthermore, PTP1B deletion in microglia results in elevated glycolytic output^37^, suggesting a connection between PTP1B activity and glycolysis that could plausibly be mediated by direct dephosphorylation of HK1. Consistent with these observations, HK1 is a known interactor of PTP1B^39^, and it was enriched by all 8 PTP1B-AbK variants, even in the absence of EGF (**Table S1**). Critically, this enrichment increased from a log_2_ fold-change +/-UV of 1.5 to a log_2_ fold-change +/-UV of 4 in response to EGFR stimulation. Notably, PTP1B was able to dephosphorylate a phosphopeptide spanning the HK1 Y732 site with similar Michaelis-Menten parameters to that of common PTP1B substrates, such as EGFR tail phosphosites (**Figure S4**)^63^. These results suggest that PTP1B dephosphorylates HK1 pY732 and could negatively regulate the cell’s ability to shift to glycolysis.

Several proteins in our +/- EGF analysis were depleted in both the L119AbK and L119AbK+D181A conditions (**Figure 6D,E**). These might be proteins that are dephosphorylated upon EGFR stimulation but still captured by photo-crosslinking. For example, RHEB and FAM83D were depleted in both conditions and are indeed directly involved in the EGFR signaling cascade^72,73^. FAM83G and FAM83B are known substrates of PTP1B, further hinting that close homolog FAM83D is likely to be a substrate^21^.

Interestingly, we also identified some EGF signaling-related proteins that were enriched in the EGF-stimulated L119AbK samples but depleted upon addition of the D181A substrate-trapping mutation, including mTOR, MARCKS, and the known PTP1B substrate ANXA2^74–76^. This indicates that while the substrate-trapping mutation can enhance capture of some proteins, there may be cases where it is detrimental to substrate capture, at least in the context of a photo-crosslinking experiment. To examine this more deeply we compared UV-exposed L119AbK samples +/- the D181A mutation (**Table S7**). Both with and without EGFR stimulation, we observed a large number of proteins that were statistically significantly enriched or depleted upon addition of the D181A mutation on top of photo-crosslinking (**Figure 6F,G**). For example, EGFR and STAT3 capture is improved by the D181A mutation, but crosslinking of the known substrates DBN1, ATXN2L, and ZC3H18 is diminished by the D181A mutation. This suggests that, in the context of photo-crosslinking, substrate trapping enhances the capture of some proteins but may also disrupt capture for other proteins. The precise molecular mechanism for this is not known, but perhaps the D181A mutation biases substrate specificity, or it leads to sequestration of PTP1B with high-affinity or high-abundance substrates, reducing PTP1B interactions with more transient substrates.

## Discussion

Here, we have developed a method to covalently capture substrates of tyrosine phosphatases *in situ*. First, we identified eight sites to incorporate the photo-crosslinking amino acid AbK by amber codon suppression without dramatically disrupting catalytic activity. We then compared photo-crosslinking of full-length PTP1B and a truncated construct that does not anchor to the endoplasmic reticulum and showed that localization dependent changes in PTP1B interactions can be detected using our method. Many previously unknown substrate proteins were identified, most notably the ER-resident ESYT proteins that mediate ER-PM contacts. Next, we showed that site-specific photo-crosslinking can be used to capture potential PTP1B substrates in an EGF-dependent manner. Finally, we demonstrate that the combination of photo-crosslinking and substrate-trapping mutations can enhance substrate capture with some proteins but not others.

A key biological finding from our study is the identification of ER-PM tethering protein ESYT1 as a substrate of PTP1B and determination the phosphosite on ESYT1 that PTP1B regulates. There is still little known about the functional consequences of phosphorylation at Y822 on the C2D domain of ESYT1. Although phosphorylation of this site very likely impacts PIP2 binding to the C2D domain, C2C and C2E appear to be the main domains responsible for the pinching function of ESYT1^56^. Thus, further studies are needed to determine the function of the C2D domain and consequences of Y822 phosphorylation. For the related protein ESYT2, the site of dephosphorylation by PTP1B remains to be definitively validated in cells. However, PTP1B likely dephosphorylates ESYT2 pY824, based on the homology between ESYT1 and ESYT2, the observed dephosphorylation of this specific phosphopeptide *in vitro*, and the high-frequency of reported phosphorylation at this site the in PhosphoSitePlus database^4^. For ESYT2, binding of PIP2 in the pocket where Y824 resides dictates its ability to engage the plasma membrane, implying that regulation of pY824 by PTP1B would be functionally relevant^57^.

There are currently only a few robust methodologies to identify substrates of tyrosine phosphatases. These technologies each have their drawbacks, such as making inactivating mutations, requiring that protein complexes survive cell lysis, or fusing the phosphatase to a large proximity-labeling enzyme, which also identifies non-substrate interactors and co-localized proteins^18,21^. Nonetheless, prior methods have successfully led to the discovery of many new PTP substrates, which have driven much of our understanding of PTP signaling biology. Our goal was to add to this repertoire of tools by designing a method that complements existing approaches.

The approach described here also has limitations that should be considered when choosing an optimal method for PTP substrate identification. For example, although the sites chosen to incorporate the photo-crosslinker are minimally perturbing based on sequence and structural analysis, as well as activity measurements, these positions may still make important contacts with substrates. Thus, photo-crosslinker incorporation at these sites could affect the substrate specificity of the PTP and the scope of proteins captured. However, one way to potentially circumvent this issue is to perform photo-crosslinking with multiple variants and focus on overlapping hits from different variants, as we did in this study. Photo-crosslinking is also limited by being a stoichiometric reaction, unlike proximity labeling. Thus, we reasoned that adding a substrate-trapping mutation might boost the signal. Indeed, in the context of EGFR stimulation, substrate trapping greatly increased the signal for some photo-crosslinked proteins. Finally, we note that our photo-crosslinking platform was performed with overexpression of our variants. Ideally photo-crosslinking would be performed at near endogenous levels of PTP1B to obtain a more refined representation of endogenous substrates.

Overall, we have demonstrated that photo-crosslinking can capture substrates of PTP1B and is sensitive enough to capture localization- and signal-dependent substrates. We conducted our experiments in HEK 293 cells to develop the technology, but we envision future studies in disease-relevant cell lines to identify context-specific differences in PTP1B signaling. We also envision applying this technology to other tyrosine phosphatases. The sites identified for photo-crosslinker incorporation in our study were chosen by evaluating structural proximity to the substrate-binding pocket and tolerance to sequence variation, across the classical PTP family. Thus, our approach, and these sites in particular, could presumably be employed for other PTPs by identifying the analogous positions in these homologs. Naturally, these sites would have to be validated to ensure that they do not disrupt PTP-specific structural features, such as the interdomain auto-inhibition of SHP2^29^. Altogether, the photo-crosslinking approach described here expands the toolkit of fruitful approaches for identifying PTP substrates and deepening our understanding of PTP-driven cellular processes.

## Acknowledgements

We thank members of the Shah and Jovanovic labs for their helpful discussions throughout this project. We thank Kathrin Lang and co-workers for providing plasmids for amber codon suppression and AbK incorporation.

## Author Contributions

Conceptualization: A.C.J., N.H.S.

Data curation: A.C.J., Y.M., A.E.v.V.; M.J., N.H.S.

Formal analysis: A.C.J., Y.S.G., N.H.S.

Funding acquisition: M.J., N.H.S.

Investigation: A.C.J., Y.S.G., D.C.C, Y.M., M.L.

Methodology: A.C.J., M.L., A.E.v.V., N.H.S.

Project administration: N.H.S.

Resources: M.J., N.H.S.

Supervision: M.J., N.H.S.

Validation: A.C.J., Y.S.G.

Visualization: A.C.J., N.H.S.

Writing – original draft: A.C.J., N.H.S.

Writing – review & editing: A.C.J., Y.S.G., D.C.C., Y.M., M.L., A.E.v.V., M.J., N.H.S.

## Data availability

There are no restrictions on the use of any data generated in this study. All processed data supporting the results and conclusions are provided in the article and supplementary files. Raw proteomics data will be deposited in PRIDE.

## Supplementary Materials

The Supplementary Information file includes Materials and Methods and supplementary figures:

Figure S1. Mutational sensitivities in the SHP2 PTP domain at PTP1B crosslinker sites.
Figure S2. Analysis of background photo-crosslinking without AbK.
Figure S3. All volcano plots of photo-crosslinking PTP1B full-length AbK variants.
Figure S4. Dephosphorylation of peptides spanning putative substrate phosphosites. Supplementary references

Supplementary Tables are included as separate spreadsheet files.

Table S1. Comparison of photo-crosslinking across all full-length PTP1B constructs.
Table S2. Known PTP1B substrates and interactors.
Table S3. Proteins enriched by six or more full-length PTP1B-AbK variants but not wild-type.
Table S4. Comparison of photo-crosslinking by full-length and truncated PTP1B constructs.
Table S5. Comparison of photo-crosslinking with substrate-trapping and EGFR stimulation +/- UV.
Table S6. Comparison of photo-crosslinking with substrate-trapping and EGFR stimulation +/- EGF.
Table S7. Comparison of photo-crosslinking with EGFR stimulation +/- D181A.

