## Supplementary Information and Figures for "Identification of substrates of the protein tyrosine phosphatase PTP1B using *in situ* site-specific photo-crosslinking"

##### Table of contents

Materials and methods pg 2
Figure S1. Mutational sensitivities in the SHP2 PTP domain at PTP1B crosslinker sites. pg 8
Figure S2. Analysis of background photo-crosslinking without AbK. pg 9
Figure S3. All volcano plots of photo-crosslinking PTP1B full-length AbK variants. pg 10
Figure S4. Dephosphorylation of peptides spanning putative substrate phosphosites. pg 11
Supplementary references pg 12

##### Supplementary tables included as separate spreadsheet file

Table S1. Comparison of photo-crosslinking across all full-length PTP1B constructs.
Table S2. Known PTP1B substrates and interactors.
Table S3. Proteins enriched by six or more full-length PTP1B-AbK variants but not wild-type.
Table S4. Comparison of photo-crosslinking by full-length and truncated PTP1B constructs.
Table S5. Comparison of photo-crosslinking with substrate-trapping and EGFR stimulation +/- UV.
Table S6. Comparison of photo-crosslinking with substrate-trapping and EGFR stimulation +/- EGF.
Table S7. Comparison of photo-crosslinking with EGFR stimulation +/- D181A.

### MATERIALS AND METHODS

#### *Sequence and structure analysis to identify PTP1B crosslinker incorporation sites*

To identify residues for photo-crosslinker incorporation, we inspected 29 different PTP-substrate co-crystal structures (PDB codes: 1BZH, 1G1F, 1G1G, 4ZRT, 1EEN, 1EEO, 1PTT, 1PTU, 3I7Z, 3ZMP, 3ZMQ, 4QUM, 4RH5, 4RH9, 4RHG, 4S0G, 1FPR, 4GS0, 3D42, 3D44, 4ICZ, 4NND, 4GFU, 4GFV, 3BRH, 3OLR, 3OMH, 1YGR, and 1YGU). For each structure, residues on the PTP domain that contacted the substrate were identified. For structures of PTPs other than PTP1B, the corresponding interacting residue on PTP1B was determined from sequence and structure alignment. This analysis yielded a list of 45 interacting residues that made up the substrate-binding pocket (PTP1B numbering: 7, 11, 12, 13, 14, 15, 16, 24, 41, 42, 44, 45, 46, 47, 48, 49, 50, 51, 88, 90, 115, 116, 118, 119, 120, 121, 179, 181, 182, 183, 215, 216, 217, 218, 219, 220, 221, 251, 254, 258, 259, 262, 263, 264, and 265). Next, we downloaded the full Pfam alignment for protein-tyrosine phosphatase domain (PF00102), as of February 2021, which had 28,456 unique sequences. We calculated the frequency of all 20 amino acids and gaps at every position in the alignment, then filtered the alignment to only focus on positions that aligned with a residue in human PTP1B. Sites with high conservation, as defined by a single amino acid having a frequency greater than 0.5, sites adjacent to catalytic residues, and some glycine residues were removed, as these were all deemed likely to disrupt PTP1B function if substituted. This filtering yielded the eight sites discussed in the main text.

#### *Synthesis of AbK*

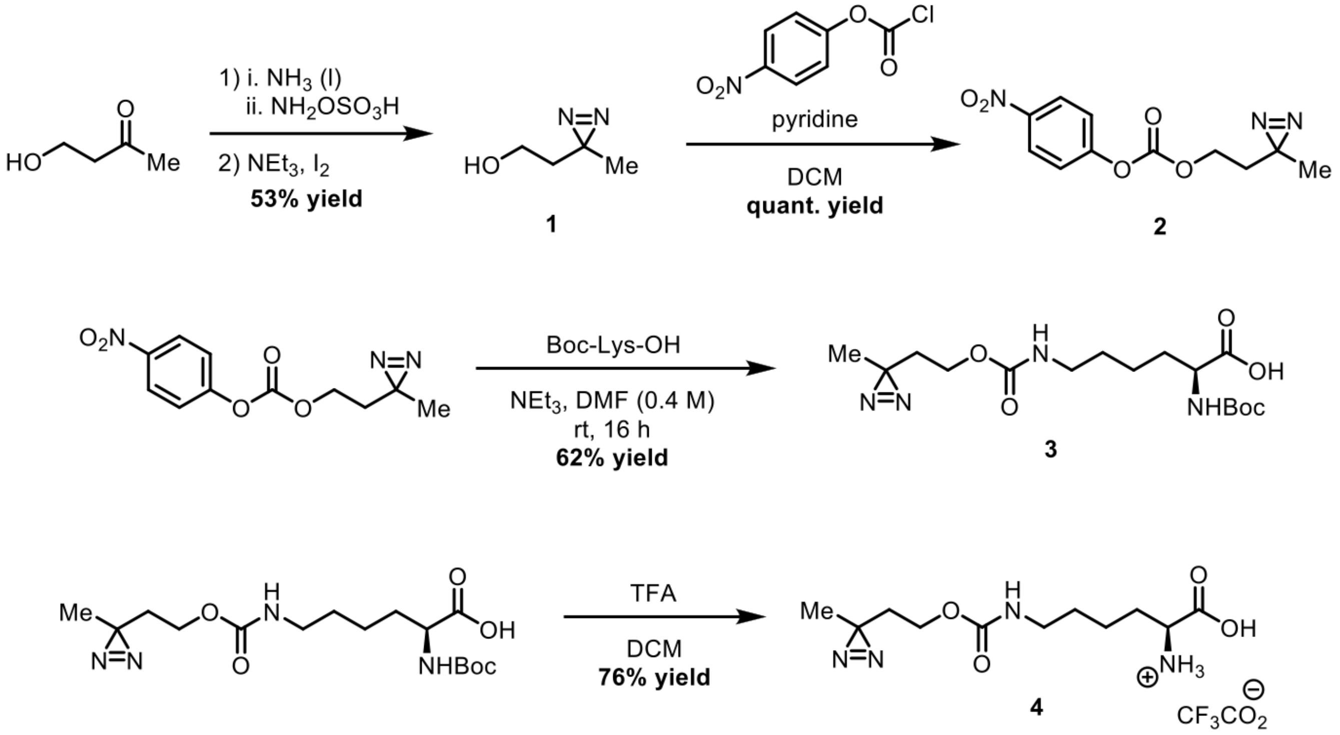


Photo-lysine was synthesized by following a previously characterized protocol and chemical shifts match that of the literature[1].

4-Hydroxy-2-butanone (1.0 g, 11.25 mmol, 1.0 eq.) was chilled on ice and $NH_3$ added (7 M in MeOH, 9 mL). After 3 hours hydroxylamine-O-sulfonic acid (1.43 g, 12.5 mmol, 1.1 eq.) was added and the reaction was stirred overnight at room temperature. The mixture was then gravity filtered and washed with MeOH. MeOH (10 mL) was then added and $Et_3N$ (2 mL) added and stirred on ice. After 3 hours the solvent was removed and $Et_2O$ (50 mL) added. The organic phase was washed with 1 M HCl (20 mL) and aqueous phase extracted again with $Et_2O$ (50 mL). The combined organic phase was washed with 20% w/v $Na_2S_2O_3$ (20 mL) and brine (20 mL). The organic phase was then dried with $Na_2SO_4$, filtered, and solvent removed with reduced pressure to yield a yellow oil. The product, **1**, did not need further purification (597 mg, 5.96 mmol, 53%).

**$^1$H NMR** (400 MHz, $CDCl_3$): δ = 1.07 (s, 3H), 1.63 (t, J = 6.3 Hz 2H), 3.53 (t J = 6.3 Hz, 2H)

DCM (11 mL) was added to a solution of **1** (200 mg, 1 mmol, 1.0 eq) on ice and 4-nitrophenyl chloroformate (242 mg, 1.2 mmol, 1.2 eq) added. Pyridine (100 µL, 1.2 mmol, 1.2 eq.) was then added and the reaction was stirred overnight at room temperature. The solvent was then removed under reduced pressure and purified by flash chromatography (10 to 15% EtOAc in hexanes) to give the final product **2** in quantitative yield (530 mg).

**$^1$H NMR** (400 MHz, $CDCl_3$): δ = 1.13 (s, 3H), 1.8 (t, J = 6.4 Hz, 2H), 4.25 (t, J = 6.4 Hz, 2H), 7.4 (d, J = 9.3 Hz, 2H), 8.29, (d, J = 9.2 Hz, 2H)

To a solution of **2** (500 mg, 1.89 mmol, 1.0 eq.) in DMF (5 mL) Na-Boc-Lysine (560 mg, 2.27 mmol, 1.2 eq.) was added with $Et_3N$ (521 µL 3.78 mmol, 2 eq.) The reaction was stirred overnight at room temperature. Heptane was then added to the mixture and was then concentrated under reduced pressure until the DMF was removed. The crude material was then purified by flash chromatography (2% MeOH in DCM to 10% MeOH in DCM) to yield the final product **3** as a colorless oil (436 mg, 1.17 mmol, 62%)

**$^1$H NMR** (400 MHz, DMSO-$d_6$): δ = 1.02 (s, 3H), 1.2-1.4 (m, 13H), 1.5-1.7 (m, 4H), 2.95 (d, J=6.4 Hz, 2H), 3.77-3.85 (m, 1H) 3.87 (t, J = 6.3 Hz, 2H), 6.98 (d, J = 7.8Hz, 1H), 7.13 (t, J = 5.6HZ, 1H)

A solution of **3** (200 mg, 0.54 mmol, 1.0 eq) in DCM (1.15 mL) was stirred on ice and TFA (496 µL) and $H_2O$ (71 µL) added. After 3 hours the solvent was removed under reduced pressure and the crude product was precipitated in ice-cold $Et_2O$. The precipitate was then collected by centrifugation and this process repeated 3 times in total. The collected precipitate was then lyophilized to obtain product **4** as a colorless powder and TFA salt (158 mg, 0.41 mmol, 76%)

**$^1$H NMR** (500 MHz, $D_2O$): δ = 1.10 (s, 3H), 1.4-1.55 (m, 2H), 1.55-1.70 (m, 2H), 1.7-1.8 (m, 2H), 1.85-2 (m, 2H), 3.2 (t, J = 6.3 Hz, 2H), 3.78 (t, J = 6.3 Hz, 1H), 4.07 (t, J = 6.0 Hz, 2H)

#### *Cloning of plasmids used for cell culture experiments*

The full-length and truncated myc-PTP1B expression vectors were reported previously[2]. The amber codon mutants were cloned using Agilent QuickChange primer design. Primers that ligate to the ends of the PTP1B construct were used with the QuickChange primers to clone the gene into two fragments containing the amber codon mutation. The two fragments were then ligated together with the insert primers to generate the full gene with the amber codon mutation. The gene was then inserted into a pEF vector backbone with Gibson assembly for mammalian expression. The substrate trapping mutation, D181A, was cloned into the gene in a similar manner. The tRNA synthetase plasmid containing 4 of the cognate tRNAs (TAN-M2) was generously gifted from Kathrin Lang's lab. The ESYT1 and ESYT2 plasmids were purchased from Addgene (plasmid #66830) and (plasmid #66831) respectively. The genes were then cloned into pEF vectors containing 3XFLAG tags using primers that anneal to the ends of the gene, with overhangs corresponding to the pEF vector. The tyrosine to phenylalanine mutation was made using QuickChange primers and a QuickChange protocol.

#### *Cell culture*

Cells were cultured in a 37°C tissue culture incubator with 5% $CO_2$. Cells were discarded by passage 25 and tested for mycoplasma every 6 months. HEK293 cells were grown in Dulbecco's Modified Eagle Medium (DMEM) with 10% Fetal Bovine Serum (FBS) and Gibco Antibiotic-Antimycotic. Throughout the methods "empty DMEM" refers to DMEM lacking FBS or Antibiotic-Antimycotic. Human epidermal growth factor was purchased in lyophilized form (#E9644, Sigma) and reconstituted in 10 mM acetic acid. Final concentrations of protease inhibitors used in lysis buffers are 0.2 mM AEBSF (4-(2-Aminoethyl)benzenesulfonyl fluoride hydrochloride), 20 µM leupeptin, and 1 µM pepstatin. Final concentrations of phosphatase inhibitor cocktail when used in lysis buffers are 2 mM activated sodium orthovanadate, 10 mM sodium fluoride, 10 mM β-glycerophosphate, and 20 mM sodium pyrophosphate.

##### *Amber suppression and photo-crosslinking test experiments*

1.5 x $10^6$ HEK 293 cells were seeded in a 6 cm plate. The next day, cells were transfected overnight with 5 µg of DNA (PTP1B-K41-Amber 1-321 truncated construct, and TAN-M2 in equal amounts) in 500 uL empty DMEM using 15 µg of polyethylenimine (PEI). Directly after transfection half of the plates received photo-lysine (AbK, 2 mM) and half did not receive AbK. The next morning, the transfection medium was replaced with pre-warmed full media (10% FBS, 1% penicillin/streptomycin). 36 h after transfection the cells were washed twice with 1 mL phosphate buffered saline (PBS) and harvested by scraping in 1 mL PBS. Cells were spun down in a refrigerated tabletop centrifuge. Cells were lysed in non-denaturing buffer (20 mM Tris-HCl, pH 8.0, 137 mM NaCl, 10% glycerol, 1% Triton-X100, 2 mM EDTA, with protease and phosphatase inhibitors added fresh) for 25 min at 4 °C while rotating, then spun down in a refrigerated tabletop centrifuge at 17,000 × *g* for 15 min. Protein concentration was determined using a bicinchoninic acid (BCA) assay. The cell lysate was then brought to 4 µg/µL. 15 µL of lysate mixture was then mixed with 15 µL of Laemmli buffer, boiled for 3 minutes, and 15 µg of total cell lysate was then loaded onto a gel. Gel was transferred onto a nitrocellulose membrane using TurboBlot (Bio-Rad), and the membrane was blocked using 5% bovine serum albumin (BSA) in Tris-buffered saline (TBS) for 1 h at room temperature. Membranes were rinsed with TBS with 0.1% Tween-20 (TBS-T) and incubated with primary antibodies in TBS-T + 5% BSA overnight at 4°C. The following primary antibodies were used, Rabbit anti-DDDDK Tag FLAG Tag, MP Biomedicals™ (purchased from Fisher #MP08L100031) 1:2500, Myc tag mouse Invitrogen (purchased from #Fisher R95025) 1:5000. Membranes were washed 5x with TBS-T and incubated with secondary antibodies (IRDye® 680RD Goat anti-Rabbit IgG, LiCor, #926-68071; and IRDye® 800CW Goat anti-Mouse IgG, LiCor, #926-32210). The blots were then washed again 5x with TBS-T Blots were imaged on a LiCor Odyssey.

The PTP1B activity assay with Src was performed similarly to the test expression. 1.5 x $10^6$ HEK 293 cells were seeded in a 6 cm plate. The next day, cells were transfected overnight with 5 µg of DNA in 500 uL empty DMEM using 15 µg of polyethylenimine (PEI). For The Src alone sample 1 µg of $Src^{Y529F}$ was transfected with 4 µg of pEF empty vector. The wild-type PTP1B sample was transfected with 1 µg of $Src^{Y529F}$, 1 µg of full-length wild-type PTP1B, and 3 µg of pEF empty vector. The same amounts were transfected for the PTP1B-D181A sample. The rest of the samples were transfected with 2 µg PTP1B-Amber construct, 2 µg TAN-M2 synthetase/tRNA plasmid, and 1 µg of $Src^{Y529F}$. Photo-lysine was added to the media directly after transfection at a final concentration of 125 µM for all samples transfected with PTP1B-Amber constructs. The samples were worked up the same as previously done for the PTP1B-AbK test expression. Two blots were produced with the first using the following primary antibodies, Myc tag mouse Invitrogen (purchased from Fisher #R95025) 1:5000, and GAPDH (14C10) Rabbit mAb (purchased from Cell Signaling Technologies #2118S). The second blot used 1:1000, Phospho-Tyrosine (P-Tyr-1000) MultiMab® Rabbit mAb mix (purchased from Cell Signaling Technologies #8954S) 1:1000. Membranes were washed 5x with TBS-T and incubated with secondary antibodies (IRDye® 680RD Goat anti-Rabbit IgG, LiCor, #926-68071; and IRDye® 800CW Goat anti-Mouse IgG, LiCor, #926-32210). The blots were then washed again 5x with TBS-T Blots were imaged on a Cytiva (Amersham) Typhoon 5 imager.

Small scale photo-crosslinking experiments were done similarly with 1.5 x $10^6$ HEK 293 cells seeded in a 6 cm plate. The next day, cells were transfected overnight with 5 µg of DNA in 500 uL empty DMEM using 15 µg of polyethylenimine (PEI). 2.5 µg of PTP1B-Amber construct were transfected with 2.5 µg of TAN-M2 synthetase/tRNA plasmid. After transfection AbK was added directly to the media with a final concentration of 125 µM. The next morning, the transfection medium was replaced with pre-warmed full media (10% FBS, 1% penicillin/streptomycin). 36 h after transfection the cells were then directly put on ice and irradiated with 365 nm light for 20 minutes using a Stratagene UV Stratalinker 2400. The samples were worked up similarly as above. The following primary antibody was used, Myc tag mouse Invitrogen (purchased from #Fisher R95025) 1:5000. Membranes were washed 5x with TBS-T and incubated with secondary antibody IRDye® 800CW Goat anti-Mouse IgG, LiCor, #926-32210). The blots were then washed again 5x with TBS-T Blots were imaged on a Cytiva (Amersham) Typhoon 5 imager

##### *Myc-tag affinity-purification mass spectrometry proteomics*

Replicates consist of separate transfections and downstream processing. $6 \times 10^6$ HEK 293 cells were seeded in a 15 cm plate. The next day, cells were transfected overnight with 25 µg DNA (Myc-tagged PTP1B-Amber, myc-tagged PTP1B-Amber-D181A, or TAN-M2) in 2.5 mL empty DMEM using 75 µg of polyethylenimine (PEI). Photo-lysine was added into the media at 125 µM final concentration directly after transfection. The next morning, the transfection medium was replaced by pre-warmed empty DMEM. 36 h after transfection the samples stimulated with EGF received empty DMEM containing 200 ng/mL epidermal growth factor (EGF) and were incubated for 10 minutes at 37 °C. Samples that were not stimulated with EGF were given empty DMEM lacking EGF and were incubated for 10 minutes at 37 °C. The cells were then directly put on ice and irradiated with 365 nm light for 20 minutes using a Stratagene UV Stratalinker 2400. After irradiation they were washed twice with 2 mL phosphate-buffered saline (PBS) and harvested by scraping in 1 mL PBS. Cells were spun down in a refrigerated tabletop centrifuge. Cells were lysed in non-denaturing lysis buffer (20 mM Tris-HCl, pH 8.0, 137 mM NaCl, 10% glycerol, 1% Triton-X100, 2 mM EDTA, with protease inhibitors added fresh) for 25 min at 4 °C while rotating, then spun down in a refrigerated tabletop centrifuge at 17,000 × *g* for 15 min. Protein concentration was determined using a bicinchoninic acid (BCA) assay. About 1500 µg of protein was used in an immunopurification using 75 µL of magnetic anti-Myc beads. Samples were left overnight at 4 °C while rotating. The next day, samples were prepared for proteomics.

##### *Preparation of proteomic IP-samples*

Samples were prepared according to a previously reported protocol with additional washes added[3]. Beads were washed four times in 1 mL 50 mM Tris-HCl (pH 8.0), then twice more in 200 µL 2 M urea in 50 mM Tris (pH 8.0). Beads were resuspended in 80 µL of 2 M urea in 50 mM Tris-HCl (pH 8.0), containing 1 mM DTT and 0.4 µg trypsin (Promega: V5113) at room temperature while shaking moderately. After 1 h, the supernatant was transferred to a new tube. Beads were washed twice with 60 µL of 2 M urea in 50 mM Tris (pH 8.0). The washes were combined with the on-bead digest 80 µL supernatant to a total volume of 200 µL. Dithiothreitol (DTT) was added to a final concentration of 4 mM, and the samples were incubated at 25 °C and shaken at 600 rpm. After 30 min, iodoacetamide (IAA) was added to a final concentration of 10 mM, and samples were incubated at 25 °C in the dark while shaking at 600 rpm. After 45 min, an additional 0.5 µg of trypsin was added to each sample. Digestion proceeded overnight at 25 °C while shaking at 600 rpm. After overnight digestion, samples were acidified using formic acid (FA) to ~1% (vol/vol). Samples were desalted using C18 StageTips (two plugs) according to Rappsilber et al.[4]. Briefly, tips were conditioned with 100 µL of 100% MeOH, 100 µL of 0.2% FA/80% Acetonitrile (ACN), and twice with 100 µL of 0.2% FA. Acidified peptides were loaded onto the StageTips and washed twice with 100 µL of 0.2% FA. Peptides were eluted with 50 µL of 0.2% FA/60% ACN, and dried in a vacuum centrifuge at room temperature. Peptides were stored at -80 °C.

##### *LC-MS/MS analysis on a Q-exactive HF for immunopurified samples*

Digested peptides from the immunopurified samples were analyzed on a Waters M-Class UPLC using a 15 cm x 75 µm IonOpticks C18 1.7 µm column coupled to a benchtop Thermo Fisher Scientific Orbitrap Q Exactive HF mass spectrometer. Peptides were separated at a 400 nL/min flow rate with a 90-minute gradient, including sample loading and column equilibration times, using solvents A (0.1% formic acid in water) and B (0.1% formic acid in acetonitrile). The detailed gradients of solvent B are below: 2% B for 1 min; linear increase to 10% B over 29 min; linear increase to 22% B over 27 min; linear increase to 30% B over 5 min; linear increase to 60% B over 4 min; linear increase to 90% B over 1 min, held for 2 min; linear decrease to 50% B over 1 min, held for 5 min; linear decrease to 2% B over 1 min; and re-equilibrated at 2% B for 14 min.

Data were acquired in data-dependent mode using Xcalibur (4.5.470.0) software; each cycle's 12 most intense peaks were selected for MS2 analysis. MS1 spectra were measured with a resolution of 120,000, an AGC target of 3e6, and a scan range from 300 to 1800 m/z. MS2 spectra were measured with a resolution of 15,000, an AGC target of 1e5, a scan range from 200–2000 m/z, and an isolation window width of 1.6 m/z.

##### *Quantification and statistical analyses of immunopurified samples*

Raw data were searched using MaxQuant (v2.0.3.0)[5] against a combined reference proteome consisting of the UniProt human proteome (UP000005640). Searches were performed using default MaxQuant parameters, with label-free quantification (LFQ) and iBAQ quantification enabled. We required 2 or more unique peptides for protein identification. Protein group intensities were normalized for the total intensity of all observable protein groups in that sample. Normalized protein group intensities were log2-transformed and averaged, from which fold-changes were calculated. *P* values were calculated using a two-tailed, heteroscedastic *t*-test.

##### *Co-immunopurification validation experiments*

About 3 × $10^6$ HEK 293 cells were seeded in a 10-cm plate. The next day, cells were transfected using 3.33 µg of each plasmid (myc-PTP1B, FLAG-ESYT1, and Src), and 30 µg PEI in 1 mL DMEM. The transfection medium was refreshed after 16 h and replaced with complete medium. After 48 h, cells were harvested by scraping in PBS. Cells were washed three times in 1 mL PBS, and lysed in 450 µL lysis buffer (20 mM Tris-HCl, pH 8.0, 137 mM NaCl, 10% glycerol, and 1% Triton-X100 + protease inhibitors) for 30 min while rotating at 4 °C. Cells were spun at 17.7 rpm for 15 min at 4 °C. Supernatant was transferred to a clean Eppendorf tube and stored at −20 °C.

Protein concentration was determined using a BCA assay, and absorbance was measured at 562 nm using a BioTek Synergy Neo2 multi-mode reader. 500 µg of protein in a total volume of 500 µL was incubated overnight while rotating at 4 °C with 25 µL of magnetic Myc-beads, for ESYT1 co-IP by PTP1B. The beads were washed 3 times on a magnetic rack using 1 mL lysis buffer. Then, 28 uL1x Laemmli buffer was added, and beads were boiled at 100 °C for 3 min. For whole cell lysates, 15 µg protein was loaded onto a gel. For IP samples, 12 µL of boiled supernatant was used. Gel was transferred onto a nitrocellulose membrane using TurboBlot (Bio-Rad), and the membrane was blocked using 5% bovine serum albumin (BSA) in Tris-buffered saline (TBS) for 1 h at room temperature. Membranes were rinsed with TBS with 0.1% Tween-20 (TBS-T) and incubated with primary antibodies in TBS-T + 5% BSA overnight at 4°C. The following primary antibodies were used, β-Actin (Cell Signaling Technologies, #4970S) 1:1000, Phosphotyrosine (Sigma Aldrich, #05-321X) 1:1000, Myc (Cell Signaling Technologies, #2278S) 1:1000, FLAG (Fisher, #MP08L100031) 1:2500. Membranes were washed 5x with TBS-T and incubated with secondary antibodies (IRDye® 680RD Goat anti-Rabbit IgG, LiCor, #926-68071; and IRDye® 800CW Goat anti-Mouse IgG, LiCor, #926-32210). The blots were then washed again 5x with TBS-T and imaged on a Cytiva (Amersham) Typhoon 5 imager.

##### *Purification of PTP1B-WT (1-321)*

pET28-His-TEV plasmid containing wild-type PTP1B with proline rich regions but lacking the ER anchoring tail (Residues 1-321) was used for purification. BL21(DE3) cells were transformed and were grown in LB supplemented with 100 µg/mL kanamycin at 37 °C until cells reached an OD600 of 0.5. IPTG (1 mM) was added to induce protein expression, which was carried out at 18 °C overnight. Cells were centrifuged and subsequently resuspended in lysis buffer (50 mM Tris, pH 7.5, 300 mM NaCl, 20 mM imidazole, 10% glycerol, and freshly added 2 mM β-mercaptoethanol). The cells were lysed using sonication (Fisherbrand Sonic Dismembrator), and spun down at 14,000 rpm for 45 min. The supernatant was applied to a 5 mL Ni-NTA column (Cytiva). The resin was washed with ten column volumes of lysis buffer and wash buffer (50 mM Tris, pH 7.5, 50 mM NaCl, 20 mM imidazole, 10% glycerol, and freshly added 2 mM β-mercaptoethanol). The protein was eluted off the Ni-NTA column in elution buffer (50 mM Tris, pH 7.5, 50 mM NaCl, 500 mM imidazole, and 10% glycerol) and brought onto a 5 mL HiTrap Q Anion exchange column (Cytiva). The column was washed using Anion A buffer (50 mM Tris, pH 7.5, 50 mM NaCl, and 1 mM TCEP). Protein elution off the column was induced through a salt gradient between Anion A buffer and Anion B buffer (50 mM Tris pH 7.5, 1 M NaCl, 1 mM TCEP). The eluted protein was cleaved at the His6-TEV tag by the addition of 0.10 mg/mL of His6-tagged TEV protease at 4 °C overnight. This cleavage cocktail was flowed through a 2 mL Ni-NTA gravity column (Thermo Fisher) to separate the cleaved protein from uncleaved protein and TEV protease. Finally, the cleaved protein was purified by size-exclusion chromatography on a Superdex 200 16/600 gel filtration column (Cytiva) equilibrated with SEC buffer (20 mM HEPES, pH 7.5, 150 mM NaCl, and 10% glycerol). Pure fractions were pooled and concentrated, and flash frozen in liquid N2 for long-term storage at −80 °C.

#### *Synthesis and purification of phosphopeptides for activity measurements*

Phosphopeptides were synthesized using 9-fluorenylmethoxycarbonyl (Fmoc) solid-phase peptide chemistry, with the standard 20 canonical amino acids (CEM) and Fmoc-Tyr(PO(OBzl)OH)-OH for phospho-tyrosine (CEM). Peptide syntheses were done using the CEM Liberty Blue automated microwave-assisted peptide synthesizer under nitrogen atmosphere, with standard manufacturer-recommended application protocols. Peptides were synthesized on MBHA rink amide resin on a 0.1 mmol scale. Nα-Fmoc amino acids (0.2 M, 6 equiv.) were activated with diisopropylcarbodiimide (DIC, 1.0 M) and ethyl cyano(hydroxyamino)acetate (OxymaPure, 1.0 M) in dimethylformamide (DMF) prior to coupling. The coupling cycles for phosphotyrosine and the amino acid directly after it were done at 75 °C for 15s, then 90 °C for 230s. All other coupling cycles were done at 75 °C for 15s, then 90 °C for 110s. Deprotection of the Fmoc group was performed in 20% (v/v) piperidine in DMF (75 °C for 15s then 90 °C for 50s), except for the amino acid directly after the phosphotyrosine which had an additional initial deprotection (25 °C for 300s). The resin was washed (4x) with DMF following Fmoc deprotection and after Nα-Fmoc amino acid coupling. All peptides were acetylated at their N-terminus with 10% (v/v) acetic anhydride in DMF and washed 4x with DMF.

After peptide synthesis was completed, including N-terminal acetylation, the resin was washed (3x each) with dimethylformamide (DMF), dichloromethane (DCM) and methanol (MeOH), and dried under reduced pressure overnight. The peptides were cleaved and the side chain protecting groups were simultaneously deprotected in 95% (v/v) trifluoroacetic acid (TFA), 2.5% (v/v) triisopropylsilane (TIPS), and 2.5% water, in a ratio of 10 µL cleavage cocktail per mg of resin. The cleavage-resin mixture was incubated at room temperature for 90 minutes, with agitation. The cleaved peptides were precipitated in cold diethyl ether, washed in ether, pelleted, and dried under air. The peptides were redissolved in a 50% (v/v) water/acetonitrile solution and filtered from the resin.

Each crude peptide mixture was purified using reverse-phase high performance liquid chromatography (RP-HPLC) on either a semi-preparatory C18 column (Agilent, ZORBAX 300SB-C18, 9.4 x 250 mm, 5 µm) with an Agilent HPLC system (1260 Infinity II), or a preparatory C18 column (XBridge Peptide BEH C18 Prep Column, 19 x 150 mm, 5 µm) with a Waters prep-HPLC system (Prep 150 LC System). Flow rate for purification was kept at 4 mL/min (semi-preparative) or 17 mL/min (preparative) with solvents A (water, 0.1% (v/v) TFA) and B (acetonitrile, 0.1% (v/v) TFA). Peptides were generally purified over a 50 minute (semi-preparative) or 20 minute (preparative) linear gradient from solvent A to solvent B, with the specific gradient depending on the peptide sample. Peptide purity and identity were assessed by mass spectrometry (Waters Xevo G2-XS QToF) and analytical HPLC (Agilent, ZORBAX 300 SB-C18, 4.6x150mm, 5µM) at a constant flow rate of 1 mL/min over a 5-95% B gradient in 10 minutes. Pure peptides were lyophilized and redissolved in 100 mM Tris, pH 8.0 for experiments.

#### *PTP1B peptide dephosphorylation measurements*

Activity measurements using phosphopeptides were done using the EnzCheck Phosphate Assay Kit (ThermoFisher) according to the manufacturer's instructions. Reactions of 50 µL were set up in a clear polystyrene flat bottom half area 96-well plate. A substrate concentration series of 0-1000 µM was used to determine kcat and KM. Reactions were started by addition of 20 nM or 40 nM PTP1B. Absorbance at 360 nm was recorded every 3-4 seconds in a span of 2 minutes using a BioTek Synergy Neo2 multi-mode reader. In all cases, the linear region of the reaction progress curve was determined by visual inspection and fit to a line. These slopes were converted from absorbance as a function of time to product formation as a function of time using standard curves measured with inorganic phosphate. Finally, these rates were corrected for enzyme concentration used in the experiment to yield $V_0$ / [enzyme] in units of ($s^{-1}$). These corrected rates were plotted as a function of substrate concentration and fit to the Michaelis-Menten equation using non-linear regression to determine catalytic parameters. Experiments were repeated three times, and the average and standard deviation of all individual replicates are reported.

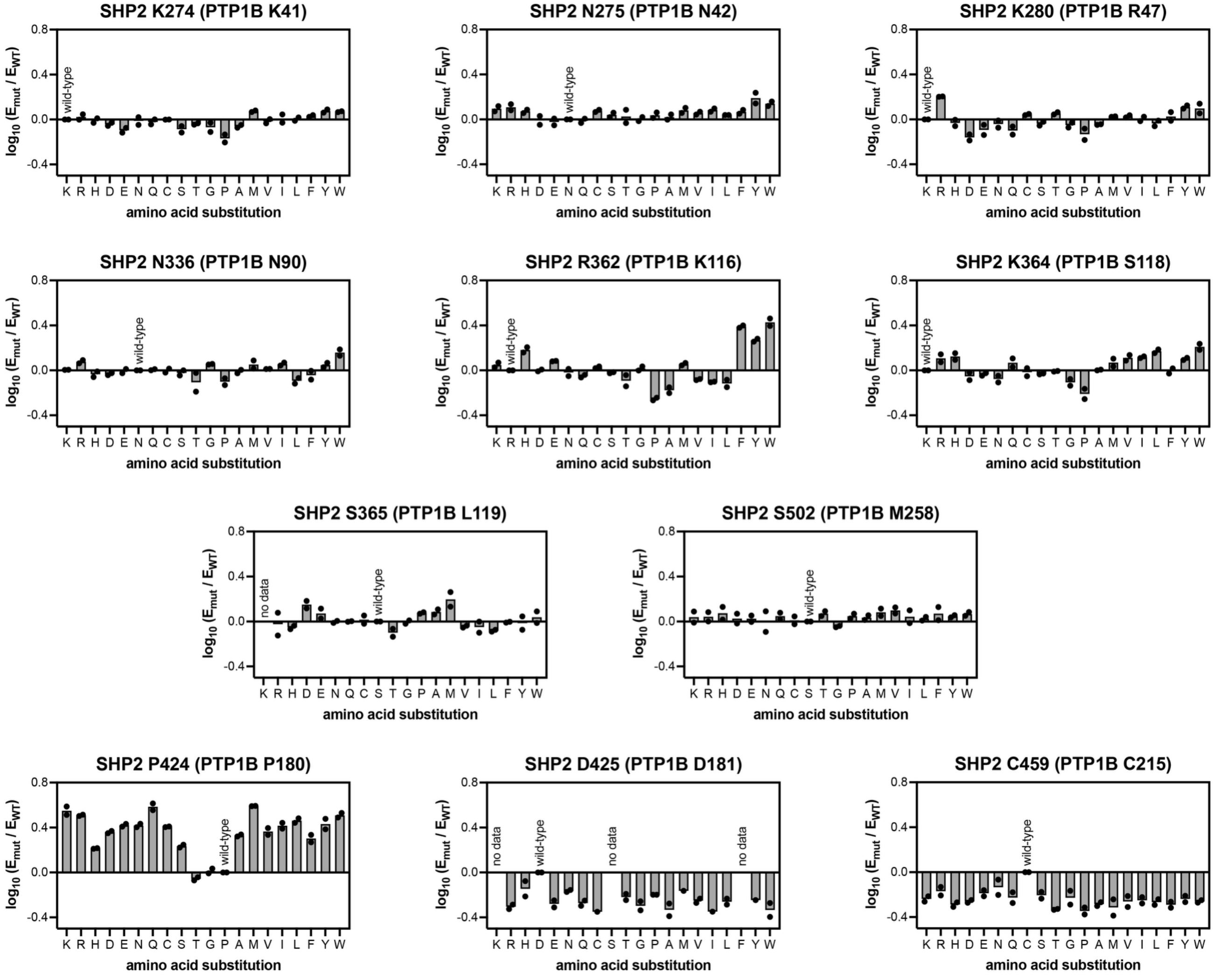


**Figure S1. Mutational sensitivities in the SHP2 PTP domain at PTP1B crosslinker sites.** Y-axes plot the $\log_{10}$-transformed enrichment for a mutant SHP2 PTP domain over the wild-type PTP domain. Enrichments are based on a previously-reported yeast selection assay, coupled with deep sequencing, in which SHP2 PTP domain variants were co-expressed with full-length v-Src, and yeast growth was dependent on SHP2 activity[6]. The top three rows show the SHP2 sites that are analogous to the PTP1B photo-crosslinker incorporation sites. For reference, the bottom row shows examples of a strongly gain-of-function mutation site (SHP2 P424) and two loss-of-function mutation sites (the catalytic cysteine, C459, and the catalytic aspartic acid, D425, which corresponds to the site of the substrate-trapping mutation in PTP1B, D181A, used throughout this study).

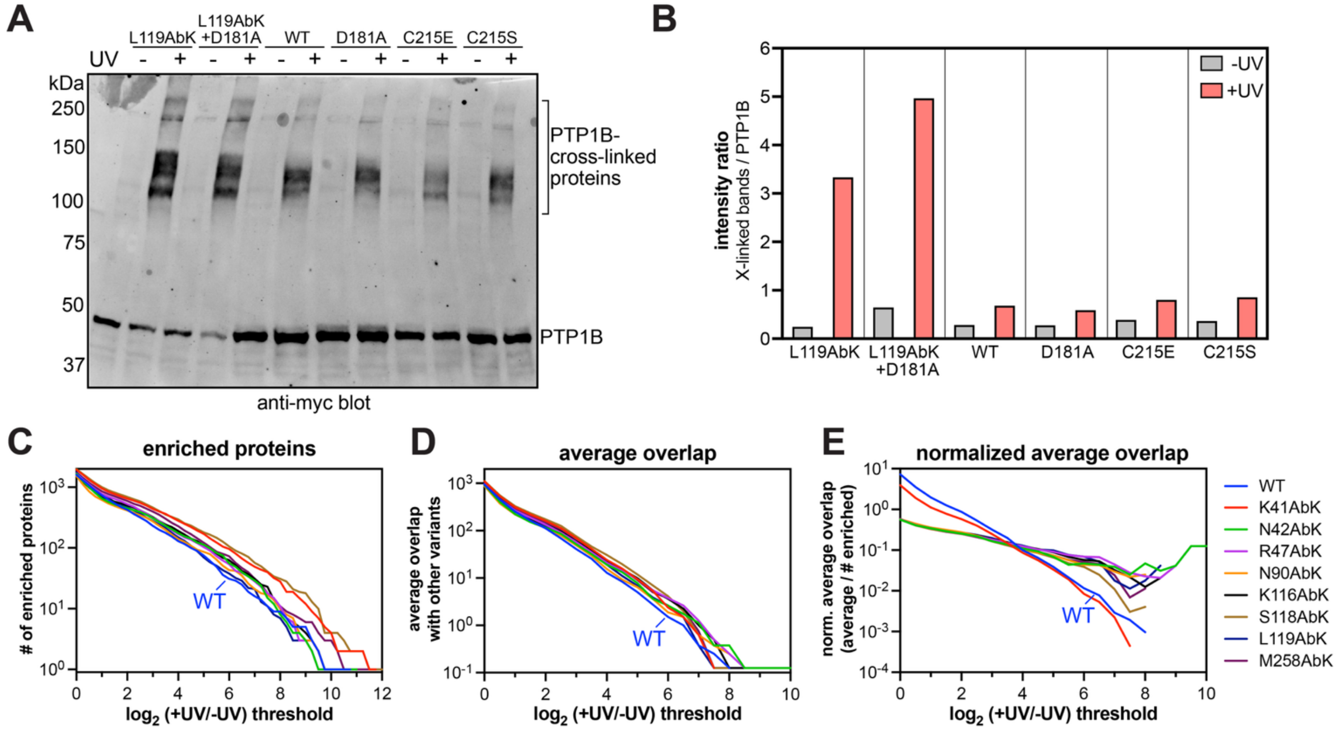


**Figure S2. Analysis of background photo-crosslinking without AbK.** (**A**) Total cell lysate anti-myc blot of PTP1B variants with or without UV irradiation. (**B**) Ratio of crosslinked band intensity to myc-PTP1B band intensity from blot in panel A. (**C**) Number of enriched proteins by wild-type PTP1B and each AbK variant, as a function of $\log_2$ fold-change for +UV/-UV comparisons, showing that wild-type has lower yields than most AbK variants at higher enrichment thresholds. (**D**) Average overlap in the number of enriched proteins between each sample and all 8 other samples at different $\log_2$ fold-change thresholds, showing that wild-type PTP1B has lower overlap with the AbK variants than most AbK variants have with each other, at higher enrichment thresholds. (**E**) Normalized overlap, calculated as the average overlap at a given fold-change threshold (panel D) adjusted for the total number of proteins enriched at that specific enrichment threshold (panel E). This graph shows that, above a $\log_2$ fold-change of 4, wild-type PTP1B and K41AbK have much lower overlap with other samples.

##### *Narrative discussion of background photo-crosslinking signal*

Most photo-crosslinking studies lack controls in which the wild-type protein is irradiated with UV light, but it is known that UV irradiation can induce some non-specific crosslinking in biomolecular systems that could confound any photo-crosslinking experiment. In particular, if the goal of site-specific photo-crosslinker incorporation is to identify site-specific interactors (e.g. proteins bound in the active site of PTP1B), nonspecific photo-crosslinking occurring elsewhere on the protein could yield false-positives when simply comparing samples with and without UV irradiation. In western blot analysis of photo-crosslinking, we consistently observed some background crosslinking by PTP1B variants lacking AbK (**Figure S2A**). To probe this further, we compared the photo-crosslinking efficiencies of wild-type PTP1B, D181A, two catalytic cysteine mutants (C215S and C215E), and two AbK variants (L119AbK and D181A+L119AbK). Both D181A and C215S are substrate-trapping mutants and should increase the lifetime of the enzyme-substrate complex relative to wild-type PTP1B. Thus, if background crosslinking is capturing substrates, we may get a boost in crosslinking signal. The C215E mutant has not been characterized in this context, but we expected a nominal impact on photo-crosslinking relative to wild-type, because it preserves the active site charge normally engendered by the catalytic thiolate. Furthermore, both C215S and C215E also test whether background PTP1B crosslinking is being mediated by the catalytic cysteine. In these comparisons, we observed UV-dependent crosslinking to PTP1B in all samples (**Figure S2A**). However, the intensities of the crosslinked bands were greater when photo-crosslinking was performed with AbK incorporated, while the intensities of bands were similar and low in the C215S, C215E, and D181A +UV samples (**Figure S2B**). These experiments suggest that proteins enriched due to background crosslinking are unlikely to be substrates but could be interactors, since they must come in close proximity to PTP1B in order to be crosslinked. To probe this further, we analyzed our proteomics data and found that wild-type PTP1B generally captures fewer proteins that AbK variants (**Figure S2C**), and wild-type PTP1B generally has less overlap in enriched proteins with other variants than each AbK variants have with each other (**Figure S2D,E**).

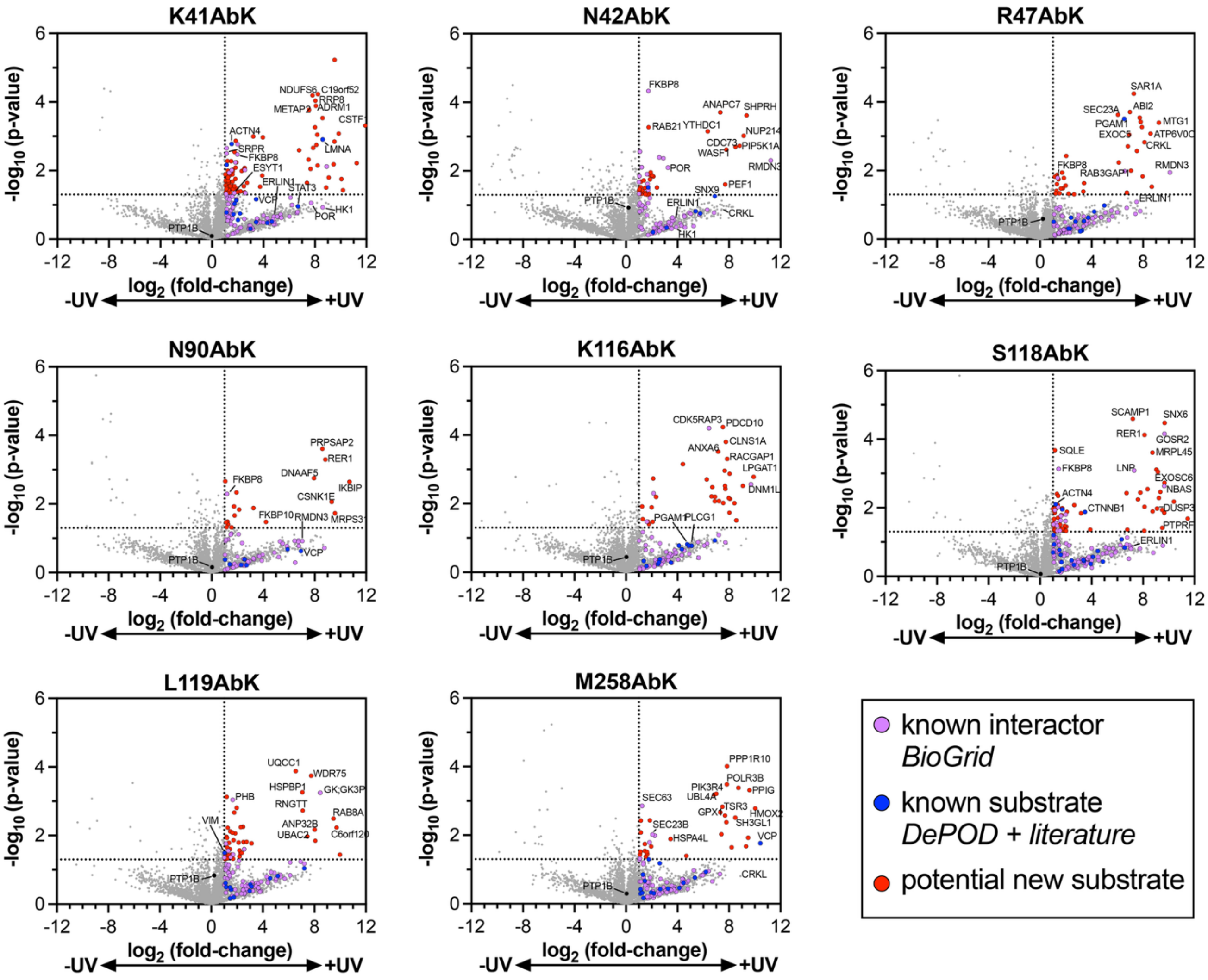

**Figure S3. Volcano plots comparing photo-crosslinking of full-length PTP1BAbK variants.** Known interactors from BioGRID are purple, known substrates from DePOD database and literature are blue, and putative substrates (proteins that are enriched >2-fold with a p-value <0.05) are in red.

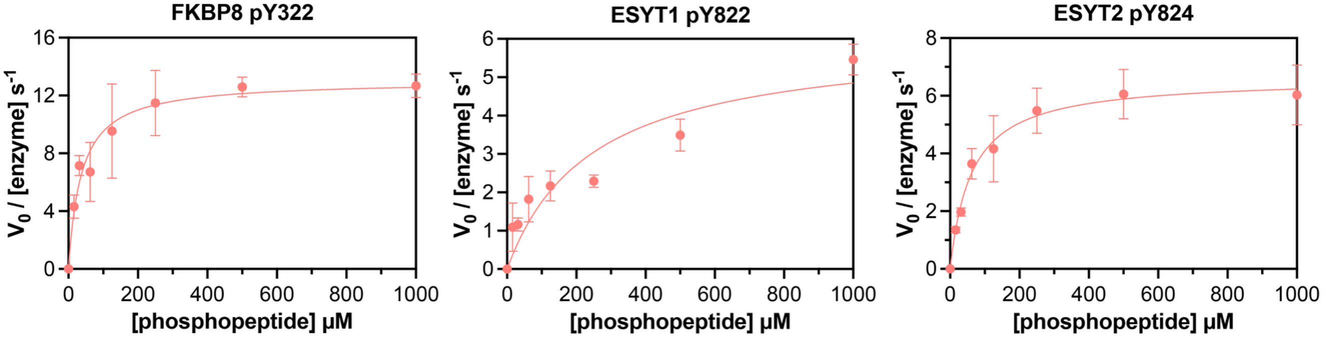


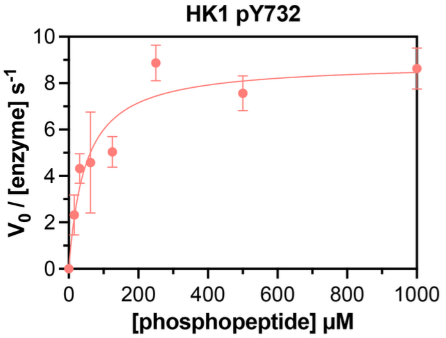


| phosphosite | peptide sequence | $k_{cat}$ ($s^{-1}$) | $K_M$ (µM) | $k_{cat}/K_M$ ($mM^{-1}\ s^{-1}$) |
|---|---|---|---|---|
| **EGFR pY1016** | `VDADE-pY-LIPQQ` | 23 ± 3 | 15 ± 5 | 1627 ± 386 |
| **EGFR pY1197** | `AENAE-pY-LRVAP` | 18 ± 2 | 15 ± 4 | 1314 ± 473 |
| **FKBP8 pY322** | `VLAQQGE-pY-SEAIPIL` | 13 ± 1 | 42 ± 14 | 353 ± 115 |
| **ESYT1 pY822** | `GTKHLSP-pY-ATLTVGD` | 6 ± 1 | 255 ± 117 | 27 ± 7 |
| **ESYT2 pY824** | `SEDGSDP-pY-VRMYLLP` | 7 ± 1 | 69 ± 25 | 108 ± 26 |
| **HK1 pY732** | `YDRLVDE-pY-SLNAGKQ` | 9 ± 1 | 53 ± 25 | 200 ± 70 |

**Figure S4. Dephosphorylation of peptides spanning putative substrate phosphosites.** (*top*) Michaelis-Menten curves for peptide dephosphorylation by PTP1B catalytic domain construct (residues 1-321). (*right*) Michaelis-Menten parameters (mean and standard deviation) for PTP1B dephosphorylation of two EGFR-derived peptides (previously reported by Chartier *et al.*[2]) and four peptides characterized in this study, all analyzed under the same assay conditions.
